# Patient and cell-type specific hiPSC-modeling of a truncating titin variant associated with atrial fibrillation

**DOI:** 10.1101/2023.03.02.530843

**Authors:** Kate Huang, Mishal Ashraf, Leili Rohani, Yinhan Luo, Ardin Sacayanan, Haojun Huang, Anne Haegert, Stanislav Volik, Funda Sar, Stéphane LeBihan, Janet Liew, Jason D. Roberts, Glen F. Tibbits, Jared M. Churko, Shubhayan Sanatani, Colin Collins, Liam R. Brunham, Zachary W. Laksman

## Abstract

**Background:** Protein truncating mutations in the titin gene are associated with increased risk of atrial fibrillation (AF). However, little is known regarding the underlying pathophysiology.

**Methods:** We identified a heterozygous titin truncating variant in a patient with unexplained early-onset AF using whole exome sequencing. We used atrial and ventricular patient induced pluripotent stem cell-derived cardiomyocytes (iPSC-CMs), CRISPR/Cas9 genetic correction, and engineered heart tissue (EHT) constructs to evaluate the impact of the titin truncating variant on electrophysiology, sarcomere structure, contractility, and gene expression.

**Results:** We generated atrial and ventricular iPSC-CMs from the AF patient with the titin truncating variant and a CRISPR/Cas9 genome corrected isogenic control. We demonstrate that the titin truncating variant increases susceptibility to pacing-induced arrhythmia (prevalence of arrhythmogenic phenotypes, 85.7% versus 14.2%; *P* = 0.03), promotes sarcomere disorganization (mean ± SEM, 66.3 ± 6.8% versus 88.0 ± 2.9%; *P* = 0.04) in atrial iPSC-CMs, and reduces contractile force (0.013 ± 0.003 mN versus 0.027 ± 0.004 mN; *P* < 0.01) in atrial EHTs compared to isogenic controls. In ventricular iPSC-CMs, this variant led to altered electrophysiology (90.0% versus 33.3%; *P* = 0.02) and sarcomere organization (62.0 ± 3.9% versus 82.9 ± 2.9%; *P* < 0.01) with no change in EHT contractility compared to isogenic controls. RNA-sequencing revealed an upregulation of cell adhesion and extracellular matrix genes in the presence of the titin truncating variant for both atrial and ventricular EHTs.

**Conclusions:** In a patient with early-onset unexplained AF and normal ventricular function, iPSC-CMs with a titin truncating variant showed structural and electrophysiological abnormalities in both atrial and ventricular preparations, while only atrial EHTs demonstrated reduced contractility. Whole transcriptome sequencing showed upregulation of genes involved in cell-cell and cell-matrix interactions in both atrial and ventricular EHTs. Together, these findings suggest titin truncating variants promote the development of AF through remodeling of atrial cardiac tissue and provide insight into the chamber-specific effects of titin truncating variants.

## Introduction

Atrial fibrillation (AF) is the most common clinical arrhythmia globally and is associated with significant morbidity and increased mortality.^1–4^ With advancements in our understanding of the genetic architecture of AF, loss-of-function truncations in the gene encoding titin (TTNtv) have been implicated in both patients with and without acquired risk factors.^5,6^ Titin is a key structural component of the sarcomere and has also been associated with dilated cardiomyopathy, arrhythmogenic cardiomyopathy, and ventricular arrhythmia.^7–9^ While recent studies of TTNtv have focused on ventricular cells, the identification of populations with unexplained AF, TTNtv, and structurally normal hearts suggests that TTNtv may have atrial specific pathophysiologic consequences.

In this study, we used atrial and ventricular patient induced pluripotent stem cell-derived cardiomyocytes (iPSC-CMs), CRISPR/Cas9 genome editing, and engineered heart tissue (EHT) to investigate the effects of an AF-associated TTNtv that was identified in a patient with early-onset AF in the absence of structural heart disease. We demonstrate that TTNtv increases susceptibility to arrhythmia, alters sarcomere organization, decreases contractility, and leads to transcriptional changes that implicate cardiac remodeling as an underlying contributor of AF pathogenesis in carriers of TTNtv.

## Methods

A detailed description of the methods is available in the Supplemental Material. Data supporting the findings of this study are available upon reasonable request to the corresponding author. This study was approved by the University of British Columbia Children’s and Women’s Research Ethics Board (H18-01465). Participants were recruited through the PERSONALIZE AF biobank (H15-02970) in Vancouver, Canada.^10–12^ All individuals provided written informed consent.

### Patient sequencing and titin variant identification

Whole exome sequencing and variant identification was performed as previously described.^12^ Briefly, genomic DNA was extracted using a Puregene DNA Blood Kit (Qiagen). Sequencing was performed using the SeqCap EZ Human Exome Library (Roche) and a NovaSeq 6000 sequencer (Illumina). Variants were filtered at a minor allele frequency of < 0.1% or missing in gnomAD. Variants with a Combined Annotation Dependent Depletion Phred score > 20 were selected and additional variant annotations were performed using dbNSFP^13^ and/or Gemini.^14^ Targeted clinical genetic testing was performed by Blueprint Genetics. Mutation carrier status were further determined through targeted Sanger genotyping (Azenta/GENEWIZ).

### iPSC reprogramming and characterization

Peripheral blood mononuclear cells were isolated from patient blood samples and reprogrammed to iPSCs using the CytoTune-iPS 2.0 Sendai Reprogramming Kit (Thermo Fisher Scientific). Genetic abnormalities were evaluated using the hPSC Genetic Analysis Kit (Stemcell Technologies) and pluripotency was assessed by immunofluorescent staining of Tra-1-60 and NANOG through confocal microscopy. Briefly, iPSCs were fixed with 4% paraformaldehyde for 20 min at room temperature, permeabilized with 0.2% Triton X-100 in DPBS without Mg^2+^/Ca^2+^ for 15 min at room temperature, and blocked overnight in 5% goat serum, 0.2% Triton X-100 in DPBS without Mg^2+^/Ca^2+^ at 4°C. Cells were stained with mouse anti-Tra-1-60 (Abcam) and rabbit anti-NANOG (Abcam) primary antibodies then incubated in goat anti-mouse AF488 (Abcam) and goat anti-rabbit AF647 (Thermo Fisher Scientific) secondary antibodies for 1 hr at room temperature. Stained coverslips were mounted on glass slides using VectaShield Antifade Mounting Media with DAPI (Vector Laboratories) and sealed with clear nail polish. A Zeiss LSM 880 Confocal Microscope with a 20X objective and Zen 2.3 SP1 FP1 version 14.0.9.201 were used to obtain maximum intensity projections of z-stack images. iPSCs were maintained in mTeSR1 media (Stemcell Technologies) on hESC-qualified Matrigel (Corning) or Geltrex (Thermo Fisher Scientific).

### CRISPR/Cas9 genome editing

CRISPR/Cas9 editing was performed using techniques adapted from published methods.^15,16^ We used CRISPR/Cas9 homology-directed repair to correct the heterozygous WT/TTNtv to WT/WT. Custom gRNA sequence design and off-target prediction were performed using CRISPOR version 4.97^17^ and the CRISPR/Cas9 gRNA Design Checker (Integrated DNA Technologies). gRNA and 127-bp DNA donor template sequences were custom ordered from Integrated DNA Technologies (Table S1). The Cas9 nuclease and gRNA duplex (dual-tracrRNA:crRNA) were delivered as a ribonucleoprotein complex with the DNA donor template using a Nucleofector 2b Device and the Human Stem Cell Nucleofector Kit 1 (Lonza) following manufacturer’s instructions. Alt-R HDR Enhancer V1 was used to improve editing efficiency (Table S2). Colonies were manually isolated, and genomic DNA were extracted using QuickExtract DNA Extraction Solution (Epicentre). PCR amplification products of the mutation site were used in restriction fragment length polymorphism assays with the AluI restriction enzyme (NEB) to screen for possible edited clones. Genotypes were confirmed through Sanger sequencing of the purified PCR products by Azenta/GENEWIZ. Off-target editing was assessed through PCR amplification and Sanger sequencing of the top five predicted sites (Table S3).

### iPSC-CM differentiation

iPSCs were differentiated into ventricular-like iPSC-CMs through temporal modulation of WNT signaling.^15,18,19^ iPSCs were treated with CHIR99021 (Stemcell Technologies) for 24 hrs (day 0-1) in RPMI 1640 with B27 minus insulin (Thermo Fisher Scientific) followed by IWP-2 (R&D Systems) for 48 hrs (day 1-3). Media was refreshed every two days until day 7 and switched to RPMI 1640 with B27 with insulin. At day 10, iPSC-CMs underwent metabolic selection using RPMI 1640 with B27 without D-glucose (Thermo Fisher Scientific) supplemented with sodium L-lactate (Sigma-Aldrich) for 4-6 days. Following metabolic selection, iPSC-CMs were maintained in RPMI with B27 media for subsequent experiments. Atrial-like iPSC-CMs were generated as described above except RPMI 1640 with B27 minus insulin media was supplemented with 0.75 μM all-trans retinoic acid (Sigma-Aldrich) for 48 hrs during days 3-5 of differentiation.^20,21^ Experiments were conducted by using iPSC-CMs > 30 days in culture.

### Optical mapping of voltage and calcium transients

iPSC-CMs were co-stained with FluoVolt (Thermo Fisher Technologies) and Calbryte 630 AM (AAT Bioquest) in RPMI 1640 media supplemented with 0.02% pluronic F-127 (Thermo Fisher Technologies) and 1.25 μM probenecid (AAT Bioquest). iPSC-CMs were imaged in a stage-top incubator at 37°C and 20 seconds recordings were acquired at 10X, 200 frames per second using a Nikon Ti2-E microscope equipped with a pco.edge 4.2Q High QE sCMOS camera. Electrical stimulation was applied using carbon Pacing Electrodes (EHT Technologies) attached to a custom-built stimulator (biphasic 5 ms pulse). The initial pacing frequency was set to a frequency closest to the spontaneous beat rate then increased stepwise by increments of 100 ms per recording. The experiment was terminated when an arrhythmogenic event was observed or until the pacing frequency reached a cycle length of 200 ms in atrial iPSC-CMs and 300 ms in ventricular iPSC-CMs, whichever came first. An arrhythmogenic event was defined as the presence of early-after depolarizations, delayed-after depolarizations, alternans, missing/extra beats, or irregular rhythms in the voltage and/or calcium transient recording. Data were processed and analyzed using a custom-built Python analysis script. Electrophysiological parameters assessed included beat rate, action potential durations (80% repolarization, 50% repolarization, and 30% repolarization), and calcium transient durations (80% repolarization, 50% repolarization, and 30% repolarization). Beat rate-correction was calculated using Bazett’s method:

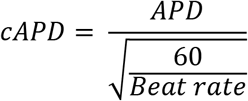

### Sarcomere organization analysis

iPSC-CMs were seeded at 30,000 cells per well on Matrigel-coated micropatterned coverslips (Curi Bio) and cultured for 21 days. iPSC-CMs were fixed, permeabilized, and blocked as described above for immunofluorescence staining of iPSCs. iPSC-CMs were stained with mouse anti-α-actinin (Sigma-Aldrich) and rabbit anti-titin M-line (Myomedix) primary antibodies overnight at 4°C then incubated in goat anti-mouse AF488 (Abcam) and goat anti-rabbit AF647 (Thermo Fisher Scientific) secondaries for 1 hr at room temperature. Stained coverslips were mounted on glass slides using VectaShield Antifade Mounting Media with DAPI and sealed with clear nail polish. Images were acquired with a 20X objective at 3X zoom using a Zeiss LSM 880 Confocal Microscope equipped with an AiryScan detector and Zen 2.3 SP1 FP1 version 14.0.9.201. Sarcomere organization was quantified using scanning gradient Fourier transform^22^ in MATLAB version R2020b.

### iPSC-CM expansion and engineered heart tissue construction

iPSC-CMs were expanded following a published protocol.^23,24^ After metabolic selection, iPSC-CMs were dissociated and plated onto Matrigel- (Corning) or Geltrex- (Thermo Fisher Scientific) coated 6-well plates at 5×10^5^ cells/well in RPMI 1640 with B27 and 10% knockout serum replacement (Thermo Fisher Scientific). Cells were expanded by culturing in RPMI 1640 with B27 with 2 μM of CHIR99021 (Stemcell Technologies). Media was replaced daily for 6-7 days or until cells reached confluency. Expanded iPSC-CMs were dissociated for subsequent EHT construction. EHTs were generated in a fibrin-based hydrogel format according to the EHT Technologies product manual. Modifications were made to the composition of the hydrogel and the EHT culture media. DMEM and horse serum were replaced by RPMI 1640 basal medium, and B27 with insulin; The hydrogel was prepared by mixing the following components on ice: RPMI 1640, B27 with insulin, GlutaMAX (Thermo Fisher Scientific), water for injection, penicillin-streptomycin, ~1 mg/mL of Matrigel and 5.4 mg/mL of fibrinogen (Sigma-Aldrich). Cell mixtures were made by combining iPSC-CMs with the hydrogel, Y-27632 (Selleck Chemicals), and thrombin (Sigma-Aldrich). A mixture of 1×10^6^ iPSC-CMs was used to construct each EHT, and EHTs were cultured in media containing RPMI 1640 with B27 supplement, 33 μg/mL of aprotinin (Sigma-Aldrich), and 50 U/mL of penicillin-streptomycin. Media changes were performed every other day and spontaneous contraction of the EHTs was observed a week post-casting.

### Contractile force measurements

EHTs were imaged in a stage-top incubator at 37°C using a 4X objective. Recordings were acquired using a Nikon Ti2-E microscope equipped with a pco.edge 4.2Q High QE sCMOS camera. A custom MATLAB-based program was used to determine the contractile force measurements by detecting tissue post deflection and converting the displacement distance into force. Background signals were removed using a low-pass filter. Using MATLAB’s peak detection function, variables such as average contraction force, contraction velocity, relaxation velocity, contraction time (T1), relaxation time (T2), and resting time were extracted from the trace signal. Average contraction force is calculated by taking the average of all the peak amplitudes. Contraction and resting velocity are derived by dividing the 80% of maximum amplitude with T1 and T2, respectively. T1 and T2 are calculated by extrapolating past half-width-maxima to determine 20% contraction. Resting time is calculated by taking the difference between two peaks at 0% contraction.

### Bulk RNA-sequencing and analysis of differential expression and splicing

Total RNA integrity was assessed with the TapeStation 4200 (Agilent Technologies) and quantified with the Qubit RNA HS assay (Thermo Fisher Scientific). 200ng of total RNA was used as input for ribosomal RNA depletion using the MGIEasy rRNA Depletion kit (MGI). Strand-specific libraries were prepared the MGIEasy RNA Directional Library Prep kit (MGI). Library quality and size were checked with the TapeStation 4200 D1000 Assay (Agilent Technologies) then quantified using the Qubit dsDNA HS Assay (Thermo Fisher Scientific). Libraries were sequenced on a MGI DNB-G50 sequencer to generate approximately 40 million 100-bp paired-end reads per library. Fastq files were aligned to the reference transcriptome with STAR version 2.6.0a, quantified with HTSEQ version 0.11.2 and normalized/differential expression was calculated with DESEQ2 version 1.16.1. Genes with a log2-fold change > 1.5 and an adjusted *P*-value < 0.05 were considered significantly differentially expressed. Differential splicing was assessed using the AS-Quant pipeline.^25^ Splicing events with FDR-adjusted *P*-values < 0.05 and absolute ratio differences > 0.1 were determined to be significantly differentially spliced. A ratio difference < 0.1 indicates a splicing event in WT/WT samples, while a ratio difference > 0.1 indicates a splicing event in WT/TTNtv samples.

### Statistical Analysis

Statistical analyses were performed using R version 4.2.1. For two group comparisons, data were analyzed using Welch’s t-test assuming unequal variances or Mann-Whitney U test. When stated, multiple testing correction was performed using the Benjamini-Hochberg procedure. Sample matching for EHT contractility analysis was performed using the coarsened exact matching method from the MatchIt R package.^26^ Matched samples were analyzed through univariate linear regression and by incorporating matching probability weights. Data plots are displayed in mean ± SEM unless otherwise specified, and *P*-values < 0.05 were determined to be statistically significant.

## Results

### Atrial fibrillation patient iPSC reprogramming and CRISPR/Cas9 genome editing

The proband was diagnosed with early-onset ‘lone’ atrial fibrillation at 18 years old in the absence of known clinical risk factors. Clinical characteristics of the patient at the time of study recruitment are described in Table S4. Notably, the patient did not exhibit signs of cardiomyopathy on cardiac MRI. We performed whole exome sequencing of participants in the PERSONALIZE AF biobank, and a heterozygous titin truncating variant (TTNtv) was identified in this patient. The TTNtv is in an exon with a proportion spliced in score > 90^27^ and located in the A-band region of titin. The American College of Medical Genetics and Genomics classification guidelines^28^ denote this variant to be likely pathogenic as it leads to a frameshift and early translation termination (Figure 1A).

**Figure 1.**
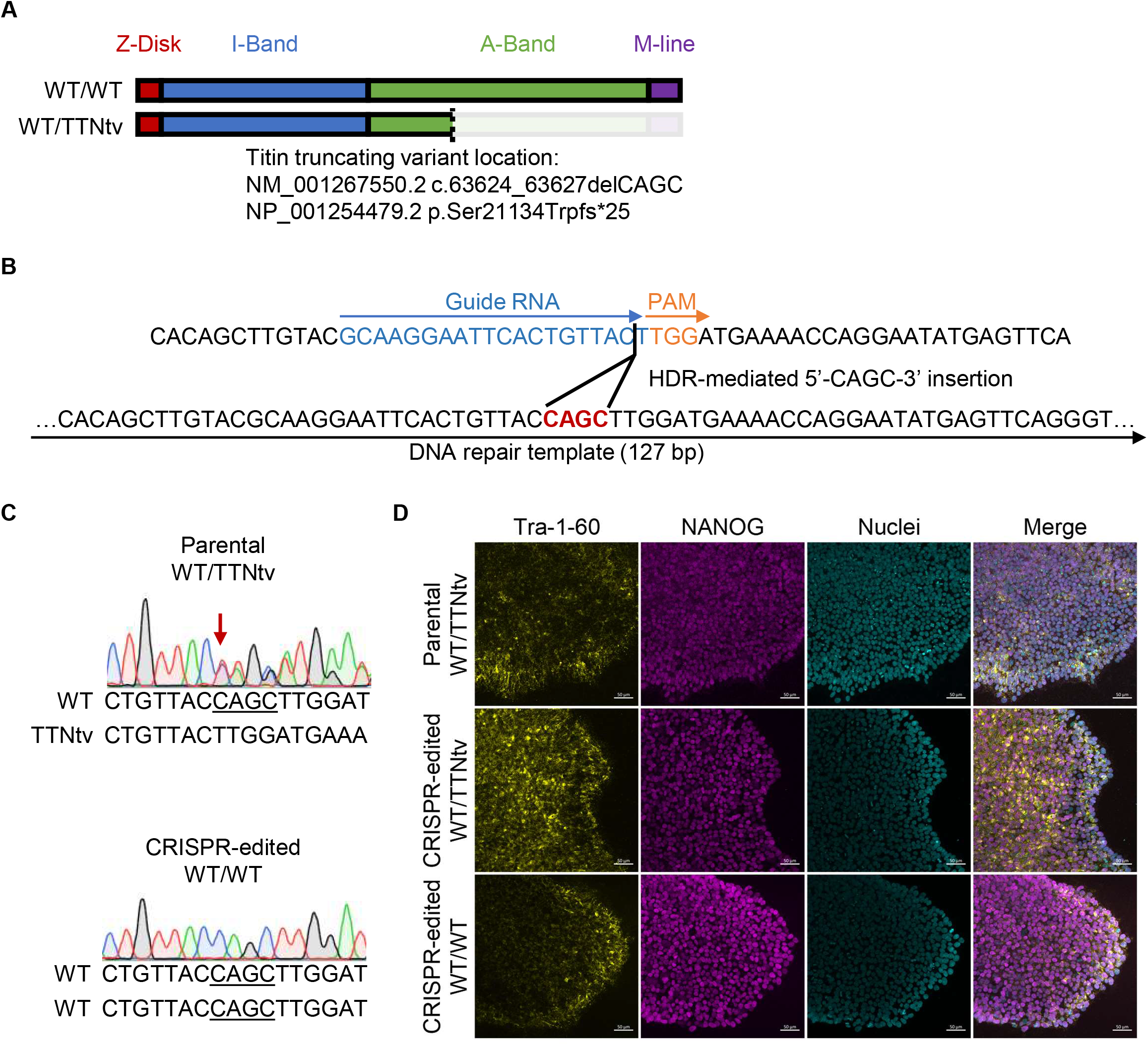
Patient iPSC reprogramming and CRISPR/Cas9 gene editing. **A**, Diagram of titin truncating variant location. **B**, Schematic of CRISPR/Cas9 gene editing approach to revert the mutation to wild-type through a 4-bp insertion. **C**, Sequence chromatograms of parental iPSCs and CRISPR-edited clones. **D**, Representative immunofluorescent images showing iPSCs expressing pluripotency markers TRA-1-60 (yellow), NANOG (purple), and DAPI (cyan). Scale bars = 50 μm. TTNtv indicates titin truncating variant; WT, wild-type.

To study the effects of the patient’s TTNtv in an *in vitro* model, we reprogrammed peripheral blood mononuclear cells into iPSCs. We used CRISPR/Cas9-mediated homology-directed repair to correct the heterozygous variant (WT/TTNtv) in the AF patient iPSCs to wildtype (WT/WT), producing a pair of isogenic iPSC lines that differ only by the presence or absence of the titin variant (Figure 1B and 1C). The iPSCs expressed the pluripotency markers TRA-1-60 and NANOG, indicating their potential for successful directed differentiation into iPSC-CMs (Figure 1D).

### Assessing TTNtv using cell type-specific iPSC-CM models

We used previously described differentiation techniques^15,18–21^ to generate atrial and ventricular isogenic iPSC-CMs to study chamber-specific effects (Figure 2A). The atrial and ventricular iPSC-CMs expressed high levels of the pan-cardiac marker cardiac troponin T (Figure 2B) and expression of the ventricular paralog of myosin regulatory light chain *(MYL2)* was significantly higher in the ventricular iPSC-CMs (Figure 2C). At the mRNA level, atrial and ventricular iPSC-CMs expressed comparable levels of the pan-cardiac gene *TNNT2* while *TTN* expression was significantly higher in ventricular iPSC-CMs (Figure 2D). iPSC-CMs also showed marked differences in the expression of known atrial and ventricular genes^20,29^, such as an upregulation of *TBX5, NPPA, KCNJ3,* and *KCNA5* in atrial iPSC-CMs and upregulation of *IRX4* and *MYL2* in ventricular iPSC-CMs (Figure 2D and S1A). To assess the electrophysiological differences between cell types, we used an optical mapping approach to measure changes in voltage and calcium transients in iPSC-CMs under spontaneous beating conditions. The iPSC-CMs exhibited distinct electrophysiological phenotypes with atrial iPSC-CMs having more triangular action potential shapes, faster spontaneous beat rates, and shorter rate-corrected action potential durations (Figure 2E and 2F). These results indicate that the differentiation protocols employed efficiently directed the WT/TTNtv and WT/WT iPSCs into atrial and ventricular cardiac lineages.

**Figure 2.**
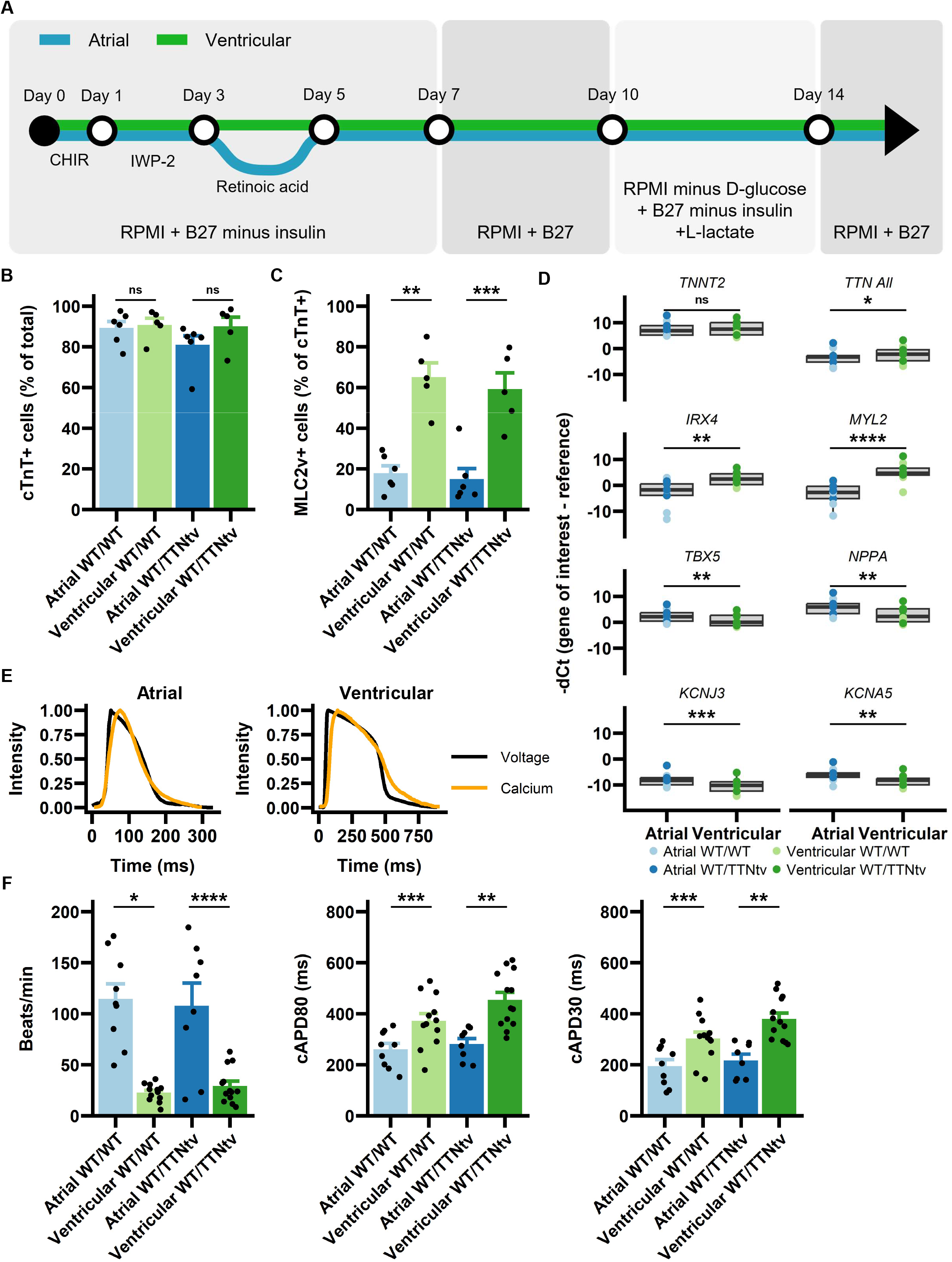
Differentiation of patient iPSCs to atrial and ventricular iPSC-CMs. **A**, iPSC-CM differentiation process. **B**, Assessment of cardiac differentiation efficiency via expression of pan-cardiac marker cTnT by flow cytometry (n = 5-6 batches). Data are presented as mean ± SEM; t-test. **C**, Expression of MLC2v in atrial and ventricular iPSC-CMs assessed by flow cytometry (n = 5-6 batches). Data are presented as mean ± SEM; t-test. **D**, RT-qPCR analysis of cardiac gene expression in atrial and ventricular iPSC-CMs (n = 4-5 batches per genotype). Data are presented as median ± IQR; whiskers represent 1.5×interquartile range; paired t-test. **E**, Representative action potential (black) and calcium transient (orange) averaged traces in atrial and ventricular iPSC-CMs. **F,** Electrophysiological characterization of atrial and ventricular iPSC-CM spontaneous beat rate, rate-corrected action potential durations at 80% repolarization and 30% repolarization (n = 8-13 batches). Data are presented as mean ± SEM; t-test. cTnT indicates cardiac Troponin T; iPSC-CM, induced pluripotent stem cell-derived cardiomyocytes; MLC2v, ventricular paralog of myosin light chain 2; \**P* < 0.05; \*\**P* < 0.01; \*\*\**P* < 0.001; \*\*\*\**P* < 0.0001.

### TTNtv increases susceptibility to pacing-induced arrhythmia in atrial and ventricular iPSC-CMs

To evaluate the effects of the TTNtv on electrophysiology, we performed optical mapping of voltage and calcium transients of spontaneously beating atrial and ventricular iPSC-CMs. We did not observe significant differences in baseline electrophysiological characteristics including beat rate (Figure S1B), rate-corrected action potential durations (Figure 3A and S1C), and rate-corrected calcium transient durations (Figure 3B and 3C) between spontaneously beating WT/TTNtv and WT/WT atrial or ventricular iPSC-CMs. We then applied electrical stimulation at increasing frequencies from a cycle length near the spontaneous beat rate to 200 ms in atrial iPSC-CMs or 300 ms in ventricular iPSC-CMs. We observed a significantly greater prevalence of arrhythmogenic phenotypes in atrial WT/TTNtv iPSC-CMs compared to isogenic WT/WT iPSC-CMs (85.7% versus 14.3%; *P* = 0.03; Figure 3D). Similarly, a greater prevalence of arrhythmogenic phenotypes were observed in ventricular WT/TTNtv iPSC-CMs compared to isogenic WT/WT iPSC-CMs (90.0% versus 33.3%; *P* = 0.02; Figure 3E). The most common arrhythmogenic phenotype observed was the appearance of voltage and/or calcium alternans at pacing cycle lengths of 200 ms or 300 ms in atrial WT/TTNtv iPSC-CMs and between 300 ms to 800 ms in ventricular WT/TTNtv iPSC-CMs (Figure 3F and 3G).

**Figure 3.**
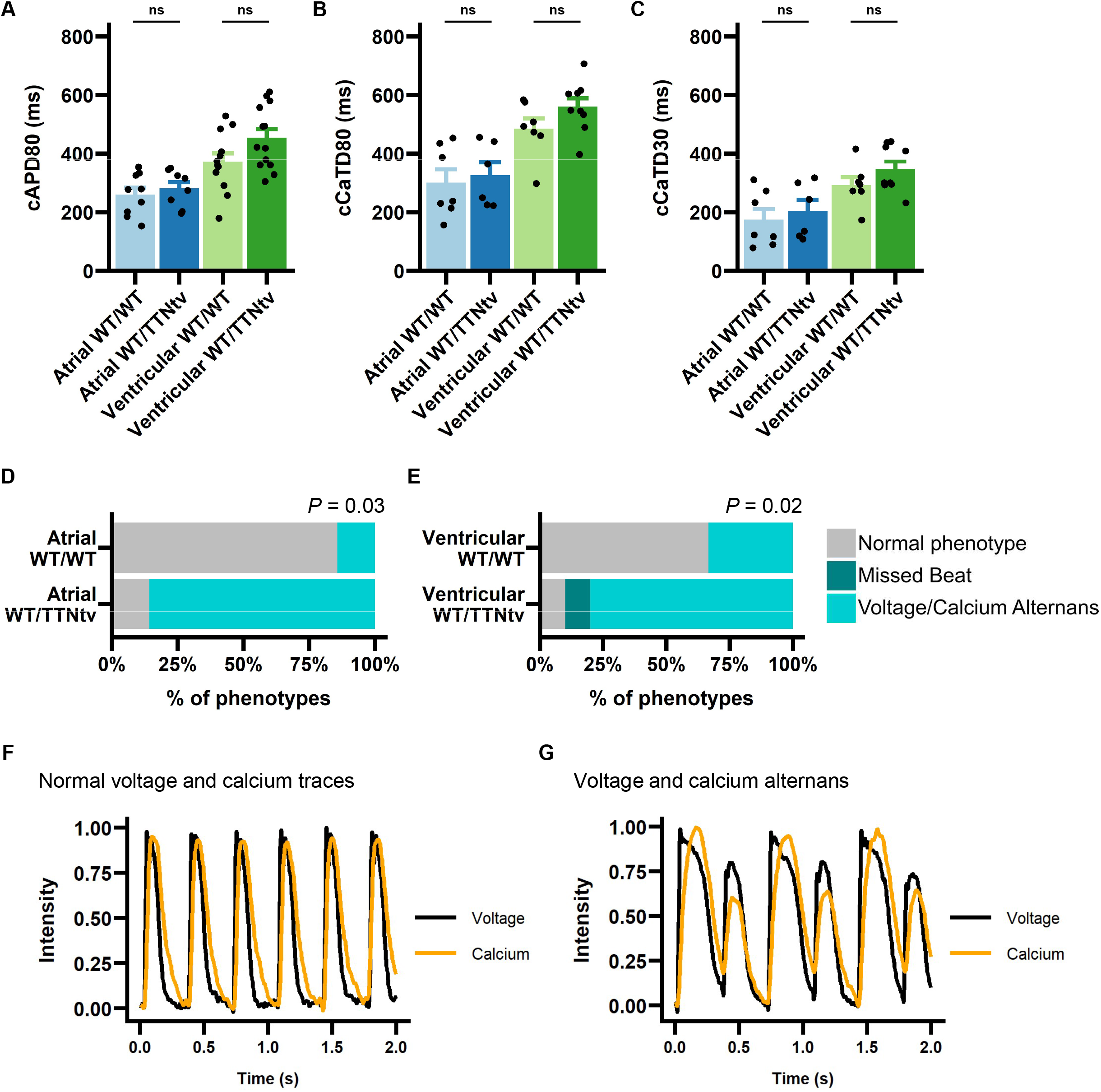
TTNtv increases the susceptibility of iPSC-CMs to pacing-induced arrhythmia. Comparisons of **A**, rate-corrected action potential duration at 80% repolarization (n = 8-13 batches), **B**, rate-correct calcium transient durations at 80% repolarization (n = 6-9 batches), and **C**, rate-corrected calcium transient durations at 30% repolarization (n = 6-9 batches) in WT/TTNtv iPSC-CMs versus isogenic WT/WT controls. Data are presented as mean ± SEM; t-test. Response of **D,** atrial iPSC-CMs (n = 7 batches) and **E,** ventricular iPSC-CMs (n = 9-10 batches) to electrical pacing for 20 seconds per cycle length. *P*-values were determined by Fisher exact test. Representative traces of iPSC-CMs with **F**, normal electrophysiology or **G**, voltage and calcium transient alternans (iPSC-CMs were co-stained with FluoVolt and Calbryte 630 AM and paced at a cycle length of 300 ms for 20 seconds). iPSC-CMs indicates induced pluripotent stem cell-derived cardiomyocytes; TTNtv, titin truncating variant; WT, wild-type.

### TTNtv impairs sarcomere structures in atrial and ventricular iPSC-CMs

Titin mediates sarcomere formation in cardiomyocytes, and previous studies have shown TTNtv can lead to sarcomere disarray or insufficiency in ventricular iPSC-CM models of TTNtv dilated cardiomyopathy.^30,31^ To determine if a TTNtv associated with AF causes sarcomere disarray in atrial iPSC-CMs, we quantified the organization of sarcomere structures in cells plated on micropatterned coverslips that are intended to promote maturation and sarcomere alignment.^32^ We observed significantly less organized sarcomere structures in atrial WT/TTNtv iPSC-CMs compared to CRISPR-corrected isogenic iPSC-CMs (mean ± SEM, 66.3 ± 6.8% versus 88.0 ± 2.9%; *P* = 0.04; Figure 4A and 4B). Ventricular WT/TTNtv iPSC-CMs also displayed disorganized sarcomere structures compared to isogenic controls (62.0 ± 3.9% versus 82.9 ± 2.9%; *P* = 0.003; Figure 4C and 4D). Taken together, the TTNtv leads to sarcomere disarray in both atrial and ventricular cell types, and genetic correction of the TTNtv can restore sarcomere structures to levels comparable to healthy controls.

**Figure 4.**
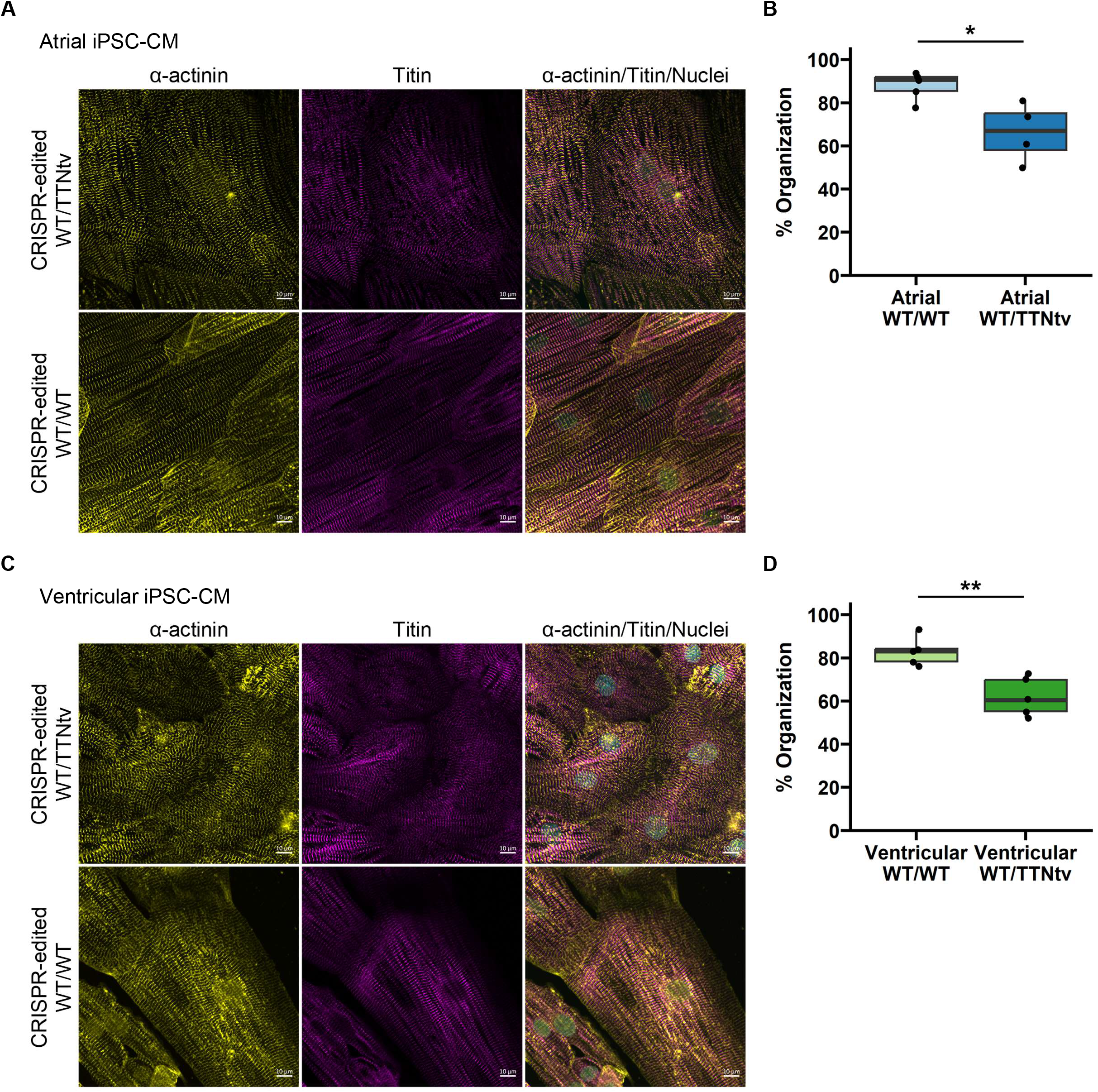
TTNtv iPSC-CMs exhibited disorganized sarcomere structures. **A**, Confocal images of atrial iPSC-CMs stained for sarcomere proteins α-actinin and titin. **B**, Quantification of sarcomere organization in atrial iPSC-CMs using sarcomere α-actinin striation patterns (n = 4-5 batches, 4-6 images each). **C**, Confocal images of ventricular iPSC-CMs stained for sarcomere proteins α-actinin and titin. **D**, Quantification of sarcomere organization in ventricular iPSC-CMs using sarcomere α-actinin striation patterns (n = 5 batches, 4-7 images each). Scale bars = 10 μm. Data are presented as median ± IQR; whiskers represent 1.5×interquartile range. iPSC-CMs indicates induced pluripotent stem cell-derived cardiomyocytes; TTNtv, titin truncating variant; WT, wild-type; \**P* < 0.05; \*\**P* < 0.01; t-test.

### TTNtv impairs contractile force generation in atrial and ventricular EHTs

We next examined the consequences of TTNtv and sarcomere disorganization on the contractile properties of atrial and ventricular iPSC-CMs. To measure contractility, we generated engineered heart tissue (EHT) constructs using atrial or ventricular iPSC-CMs (Figure 5A, 5B, and S2A). We used an imaging approach to measure force parameters based on the displacement of polydimethylsiloxane (PDMS) posts shortened by concentric contraction of the EHTs (Figure S2B). In spontaneously beating atrial EHTs, WT/TTNtv EHTs exhibited significantly weaker maximal contractile force compared to WT/WT EHTs (mean ± SEM, 0.013 ± 0.003 mN versus 0.027 ± 0.004 mN; *P* < 0.01; Figure 5C), shorter contraction time T1 (0.076 ± 0.007 s versus 0.12 ± 0.017 s; *P* < 0.05), and shorter relaxation time T2 (0.108 ± 0.02 s versus 0.181 ± 0.013 s; *P* < 0.05; Figure S2C). No significant differences were observed in contraction and relaxation velocities (Figure S2D). In ventricular EHTs, no significant differences in force, contraction and relaxation velocities, or contraction and relaxation times were observed between WT/TTNtv and WT/WT EHTs (Figure 5D, S2E, and S2F).

**Figure 5.**
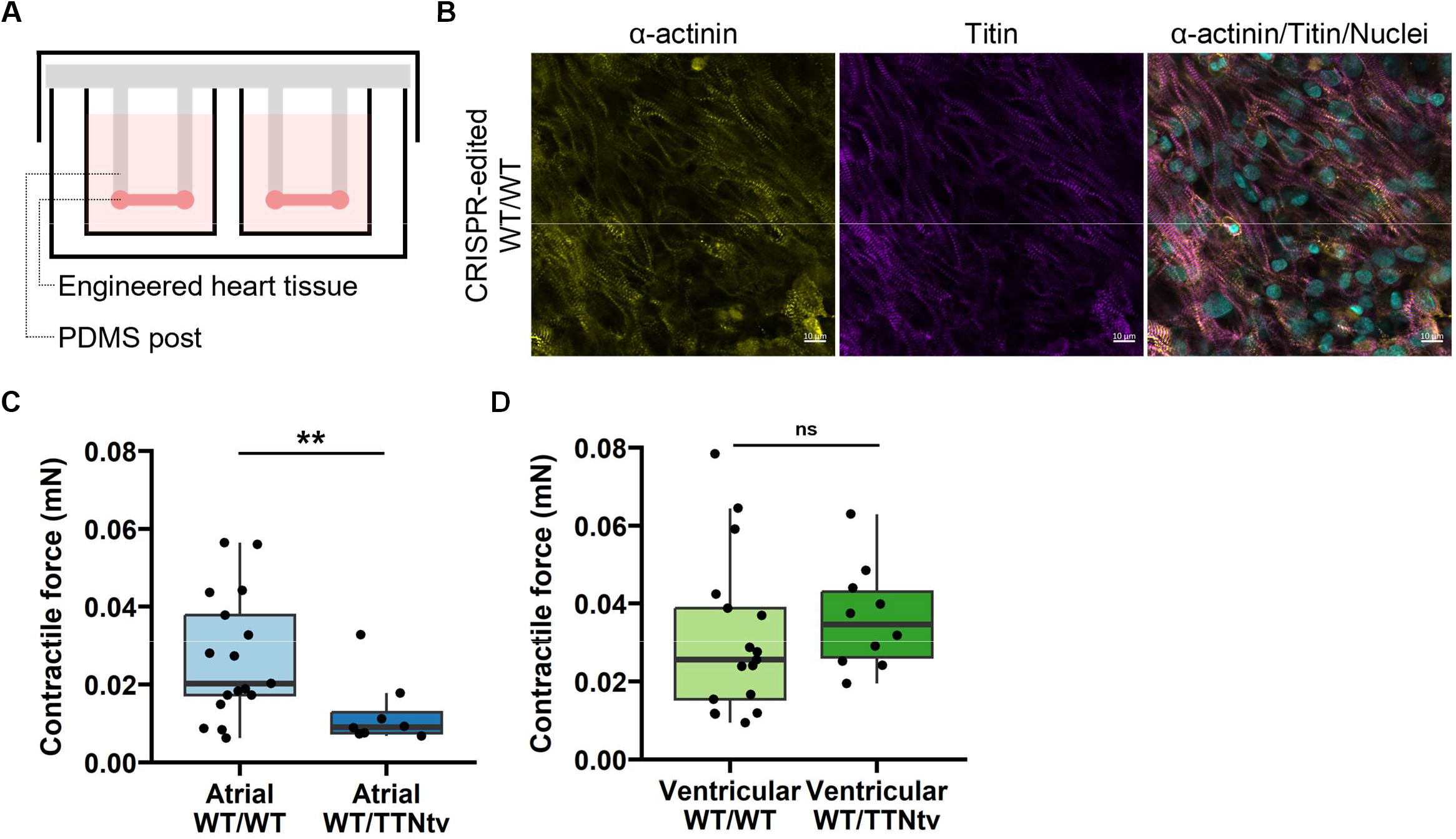
TTNtv EHTs displayed reduced contractile force compared to isogenic WT iPSC-CMs. **A**, Diagram of EHT setup. **B**, Whole-mount immunostaining of ventricular WT/WT EHTs for sarcomere α-actinin (yellow), titin M-line (purple), and nuclei (cyan). Scale bar = 10 μm. Assessment of spontaneous **C**, maximum contractile force in atrial EHTs (n = 8-17 EHTs) and, **D**, maximum contractile force in ventricular EHTs (n = 10-17 EHTs). t-test. Data are presented as median ± IQR; whiskers represent 1.5×interquartile range. EHT indicates engineered heart tissue; PDMS, polydimethylsiloxane; TTNtv, titin truncating variant; WT, wild-type; \**P* < 0.05; \*\**P* < 0.01.

To control for the effects of differing spontaneous beat rates on contractility parameters (Figure S2G), we performed a sensitivity analysis on a subset of WT/TTNtv and WT/WT EHTs matched by beat rates. In matched atrial EHTs, the contractile force of WT/TTNtv EHTs remained significantly weaker compared to WT/WT EHTs (*P* < 0.001; Table S5). Contraction velocity (*P* < 0.01) and relaxation velocity (*P* < 0.05) were also significantly slower in WT/TTNtv compared to WT/WT EHTs. No significant differences in contraction or relaxation times were detected in the matched atrial EHTs. In matched ventricular EHTs, contractile properties were not significantly different between genotypes (Table S5).

### Whole transcriptome sequencing reveals upregulation of genes involved in cell adhesion and extracellular matrix in TTNtv EHTs as well as differential splicing of cardiac-specific genes

To further understand how the TTNtv influences global gene expression in atrial and ventricular tissue, we performed whole transcriptome sequencing of atrial EHTs and ventricular EHTs (Figure 6A). *TTN* transcript expression was not significantly different between WT/TTNtv and WT/WT genotypes, similar to existing studies on *TTN* expression in heart samples of cardiomyopathy patients.^33^ We performed differential expression analysis to identify genes perturbed by the presence of the WT/TTNtv (Figure 6B). The TTNtv led to a significant downregulation of 13 genes in the atrial WT/TTNtv EHTs compared to the isogenic WT/WT EHTs (Figure 6C and Table S6), including *KLHL40,* a negative regulator of Leiomodin-3 ubiquitination that is associated with skeletal myopathies^34^ and retinoic acid receptor transcriptional regulator *RXRG* (retinoid X receptor γ subunit). Significantly downregulated genes were not detected in the ventricular samples (Figure 6D).

**Figure 6.**
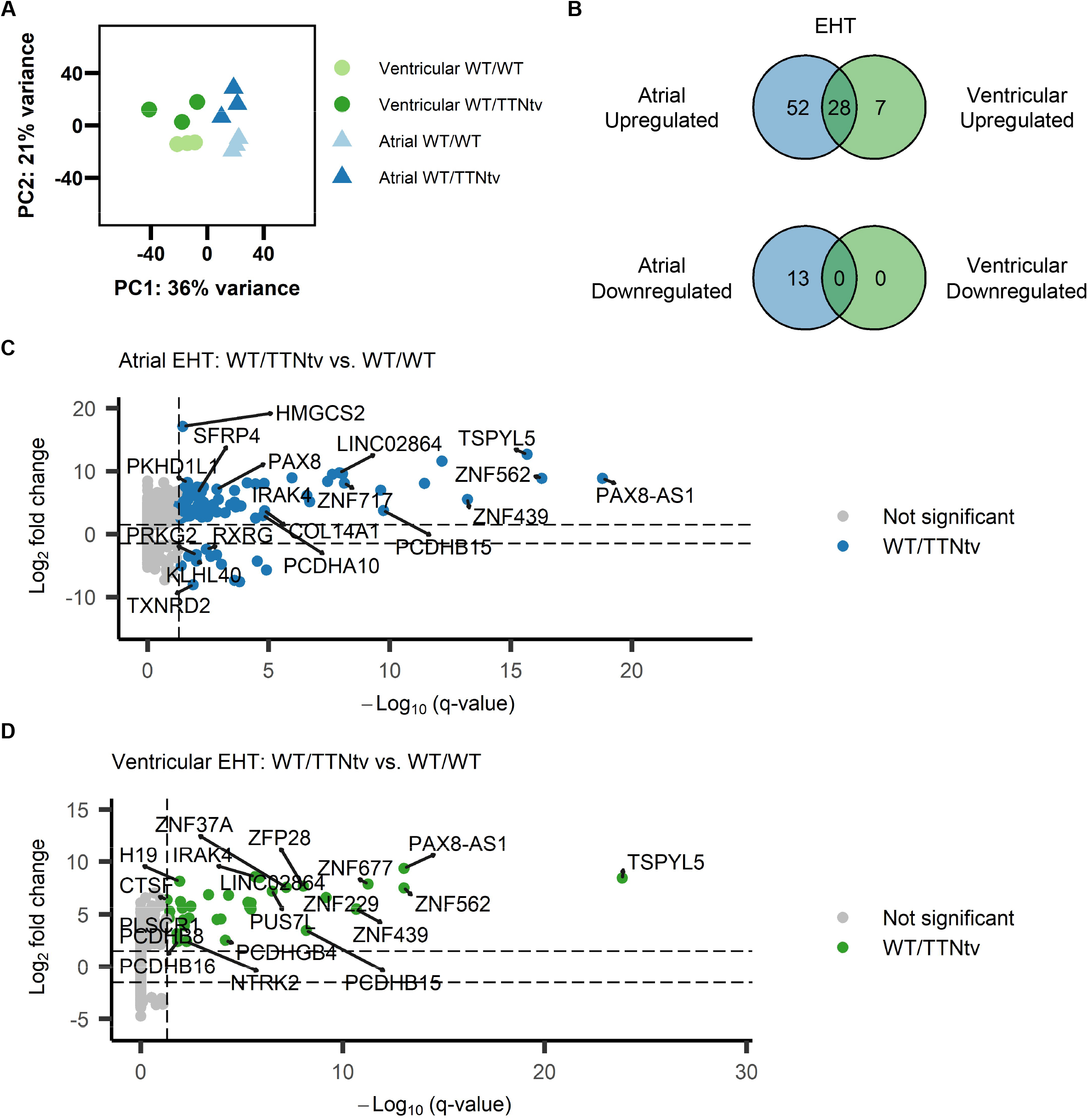
Whole transcriptome differential gene expression of atrial and ventricular EHTs. **A**, Principal component analysis of engineered heart tissue samples (n = 3 per cell type; n = 3 per genotype). **B**, Overlapping significantly upregulated and downregulated genes in atrial WT/TTNtv and ventricular WT/TTNtv EHTs relative to atrial WT/WT and ventricular WT/WT EHTs, respectively. **C**, Volcano plot of differentially expressed genes in atrial WT/TTNtv EHTs relative to isogenic WT/WT controls (blue dots indicate significant differential expression with adjusted *P*-value < 0.05 and fold change > 1.5). **D**, Volcano plot of differentially expressed genes in ventricular WT/TTNtv EHTs relative to isogenic WT/WT controls (green dots indicate significant differential expression with adjusted *P*-value < 0.05 and fold change > 1.5). EHT indicates engineered heart tissue; TTNtv, titin truncating variant; WT, wild-type.

A number of genes were significantly upregulated in the WT/TTNtv samples for both cell types compared to their respective isogenic WT/WT controls (atrial: 80 genes upregulated; ventricular: 35 genes upregulated; overlap: 28 genes; Figure 6B, 6C, 6D, Table S7, and S8). We performed over-representation analysis of the significantly upregulated genes to identify potential processes that are implicated. A significant enrichment of genes involved in cell-to-cell adhesion (GO:0007156, -log(q-value) = 2.83; GO:0098742, -log(q-value) = 1.61) as well as antiviral response (hsa05168, -log(q-value) = 5.49) were identified to be upregulated in atrial EHTs. Expression of cell adhesion *(PCDH* genes) and extracellular matrix *(COL14A1, ITGA8, LAMA3, ADAM33, SMOC2)* associated genes were also significantly upregulated, suggesting remodeling of cell-cell interactions that could have an impact on electrophysiology and contractile function. Other significantly upregulated genes identified included sodium ion channel genes *(SCN3B* and *SCN9A),* which have been linked to early-onset AF and ventricular arrhythmias.^35,36^ We also detected heightened expression of *IRAK4,* a pro-inflammatory gene which promotes cardiac remodeling post-infarction^37^ and exacerbates inflammation in myocarditis.^38^ Expression of LAMP3, which may contribute to cardiac remodeling in dilated cardiomyopathy hearts, was also significantly upregulated.^39^

We next investigated the effects of the TTNtv on alternative splicing. Splicing factors such as RBM20 and RBM24 regulate alternative splicing of several genes, including titin, that are critical to normal cardiac development and sarcomere assembly^40,41^ and have been linked to arrhythmias and cardiomyopathies when dysregulated.^42–45^ We performed differential splicing analysis of the atrial and ventricular EHT transcripts and identified significant splicing events for both atrial EHTs (Figure S3 and Table S9) and ventricular EHTs (Figure S4 and Table S10). A number of these splicing events were detected in RBM20 splicing targets. However, we did not observe significant splicing events in *TTN* transcript isoforms. Differentially spliced transcripts include those associated with the contractile apparatus *(TPM1, TPM4, TNNT2, MYL7, RBM24),* cytoskeletal genes *(ACTG1, ACTB, POSTN, ACTN1),* and adhesion or extracellular matrix genes, such as *LAMA2, EMC10,* and *COL16A1,* potentially altering protein function, domains, or protein-protein interactions and contributing to the disease phenotype. Together, these findings suggest an effect of the TTNtv on gene expression and splicing which may have an impact on atrial electrophysiology and contractile function.

## Discussion

Titin truncating variants (TTNtv) are found in approximately 0.6% of the general population, and genetic association studies have linked pathogenic titin mutations to cardiomyopathies and arrhythmias.^5,6,8,9^ TTNtv have also been linked to an increased propensity to AF in patients with and without AF risk factors.^5,6,46,47^ Existing studies to date have primarily focused on the effects of TTNtv in ventricular tissue.^31,33,48,49^ Thus, the mechanisms by which TTNtv contribute to atrial arrhythmias, particularly in the absence of ventricular disease or cardiomyopathy, remain unclear. We employed human iPSC-CMs, EHTs, and CRISPR/Cas9 genome editing to investigate the role of TTNtv in AF pathogenesis in a patient without cardiomyopathy.

We found that WT/TTNtv iPSC-CMs exhibit comparable electrophysiology to isogenic WT/WT controls at baseline. However, WT/TTNtv iPSC-CMs were significantly more susceptible to developing arrhythmogenic phenotypes upon rapid pacing with electrical stimulation. The development of AF in individuals harboring an arrhythmia-associated genetic variant may depend on the presence of additional factors or triggers^50,51^, and we have previously demonstrated an additive effect between modifiable factors, such as hypertension, and titin variant status on the risk of incident AF.^52^ Interestingly, both the atrial and ventricular iPSCs demonstrated arrhythmogenic phenotypes with rapid pacing, suggesting potentially common mechanisms and also raising the possibility that patients with AF may also be at risk of ventricular arrhythmias.

In line with titin’s known function as a critical sarcomeric structural protein^30^, we found that the TTNtv led to sarcomere disorganization in atrial iPSC-CMs. We also observed a reduction in contractile force generated by atrial WT/TTNtv EHTs, suggesting the possibility of a propensity towards an atrial-specific myopathy. As the proband was in AF for many years, an accurate assessment of atrial contractile function could not be performed. Interestingly, contractility was not impaired in ventricular WT/TTNtv EHTs which have been demonstrated previously in patient-specific models of dilated cardiomyopathy.^31,49^ While we cannot rule out the presence of subclinical structural alterations in the ventricles that were not detectable by clinical testing methods, the absence of contractility defects in the ventricular EHTs agrees with the patient’s cardiac MRI results which showed normal biventricular size and function. The TTNtv in this case demonstrated cell type-specific effects which correlate with the observed clinical phenotype and raises the possibility that unique variants in TTNtv may have varying pathophysiologic consequences that are chamber specific.

Titin mRNA was recently identified as a mediator of RBM20 nuclear foci formation which allows the splicing of multiple RBM20 target genes.^53^ We hypothesized that alternative splicing of RBM20 targets in cardiomyocytes with TTNtv may be impacted. Through whole transcriptome sequencing and differential splicing analysis of atrial and ventricular EHT transcripts, we found that titin gene expression was not significantly reduced in atrial or ventricular WT/TTNtv EHTs. Similar to a previous study on alternative splicing of titin transcripts in human dilated cardiomyopathy hearts^54^, we did not observe significant differences in the splicing of titin transcript isoforms in atrial or ventricular EHTs. We identified differential splicing of certain RBM20 targets, such as genes encoding troponin T and tropomyosin proteins, as well as transcripts not known to be regulated by RBM20, suggesting broader effects of the TTNtv on transcriptional processes in cardiomyocytes. Through differential gene expression analysis, we identified an upregulation of genes associated with cell adhesion, extracellular matrix, and inflammation, which have all been associated with AF, providing evidence to support the concept that TTNtv is involved in adverse remodeling that may contribute to arrhythmogeneisis.^55,56^ Genes encoding voltage-gated sodium channel subunits were also upregulated which could contribute to arrhythmogenesis by increasing heterogeneity of tissue refractoriness or enhancing late sodium current. Taken together, these findings may indicate that multiple and potentially patient-specific pathways and factors contribute to an increased propensity to AF. This may include both electrophysiologic derangements, as we observed, as well as an atrial-specific myopathy which would serve as substrates for AF.

This study has several limitations that should be considered. Compared to human adult heart tissue, iPSC-CMs exhibit immature electrophysiological, structural, metabolic, and transcriptional properties that more closely resemble fetal cardiomyocytes.^57^ To overcome this challenge, we used iPSC-CMs subjected to prolonged culture periods, assessed sarcomere defects in iPSC-CMs maintained on micropatterned surfaces designed to promote maturity, and generated 3D EHT constructs to compare contractility and gene expression. Further studies employing maturation techniques on EHTs or incorporating other cell types found in the heart, such as cardiac fibroblasts or endothelial cells, can improve our ability to recapitulate AF *in vitro.* While we demonstrated that the TTNtv is necessary for the development of phenotypic and functional abnormalities that are associated with AF pathogenesis, it is unclear if this specific mutation alone is sufficient to cause AF if present in other individuals. Given the varying disease phenotypes and penetrance of titin mutations in the general population, our findings do not preclude the possibility of additional factors contributing to the onset of AF in other patients with a TTNtv. More studies of this TTNtv in other genetic contexts and of other TTNtv associated with lone AF are needed to broaden the applicability of these findings.

In summary, we modeled the effects of a heterozygous TTNtv implicated in AF in atrial and ventricular patient-derived iPSC-CMs. We observed an increase in arrhythmogenicity and sarcomere disarray in atrial and ventricular TTNtv iPSC-CMs compared to isogenic controls. Using 3D EHT constructs, we showed that the TTNtv impairs contractility in atrial EHTs but not ventricular EHTs, and that the TTNtv leads to transcriptional changes which may contribute to altered cell-cell interactions and extracellular matrix composition. Our findings suggest a link between TTNtv and both structural and electrophysiological abnormalities in atrial tissue relevant to the pathogenesis of AF. These findings deepen our understanding of the effects of TTNtv on atrial and ventricular tissues and their relationship to atrial and ventricular myopathies and arrhythmias.

## Acknowledgments

The authors would like to thank the study participants who made this research possible; the BC Children’s Hospital Research Institute Tissue and Disease Modeling Core; the Centre for Heart Lung Innovation Molecular Phenotyping Core and the Cellular Imaging and Biophysics Core. RNA-sequencing was performed by the Vancouver Prostate Centre Laboratory for Advanced Genome Analysis.

## Supplementary Materials

Supplemental Methods

Tables S1–S11

Figures S1–S4

