## Supplementary Material for "Patient and cell-type specific hiPSC-modeling of a truncating titin variant associated with atrial fibrillation"

[1]Centre for Heart Lung Innovation, University of British Columbia, Vancouver, Canada.
[2]Experimental Medicine Program, University of British Columbia, Vancouver, Canada.
[3]School of Biomedical Engineering, University of British Columbia, Vancouver, Canada.
[4]Genomics Core Facility, Vancouver Prostate Centre, University of British Columbia, Vancouver, Canada.
[5]Department of Medicine, University of British Columbia, Vancouver, Canada.
[6]Population Health Research Institute, McMaster University and Hamilton Health Sciences, Hamilton, Canada.
[7]Molecular Cardiac Physiology Group, Departments of Biomedical Physiology and Kinesiology and Molecular Biology and Biochemistry, Simon Fraser University, Burnaby, Canada.
[8]Cellular and Regenerative Medicine Centre, British Columbia Children's Hospital Research Institute, Vancouver, Canada.
[9]Department of Cellular and Molecular Medicine and Sarver Molecular Cardiovascular Research Program, University of Arizona, Tucson, USA.
[10]Division of Cardiology, Department of Pediatrics, Children's Heart Centre, BC Children's Hospital, University of British Columbia, Vancouver, Canada.
[11]Vancouver Prostate Centre, University of British Columbia, Vancouver, Canada.
*Authors contributed equally

### Supplemental Methods

#### Flow cytometry

iPSC-CMs were dissociated into single cells by incubating in RPMI 1640 supplemented with 0.5U/ml Liberase TH (Sigma-Aldrich), 50U/ml DNaseI (Sigma-Aldrich), and TrypLE (Thermo Fisher Scientific) at 37°C. Cells were fixed with 2% paraformaldehyde at room temperature and permeabilized with 0.5% Saponin, 2% FBS in D-PBS. iPSC-CMs were stained with BV421-conjugated anti-cTnT antibody (BD BioSciences) and rabbit anti-MLC2v antibody (Abcam) for 1 hr then stained with an AF647-conjugated goat anti-rabbit secondary antibody (Thermo Fisher Scientific) for 1hr. iPSCs were dissociated using TrypLE at 37°C and stained in parallel to serve as a negative cell type control for gating. Stained samples were run on a Gallios Flow Cytometer (Beckman Coulter) and analyzed using the Kaluza Analysis Software version 2.1 (Beckman Coulter).

#### RT-qPCR

RNA was extracted using a Illustra RNAspin Mini Isolation Kit (Cytiva). cDNA synthesis was performed using a cDNA Synthesis Kit (Applied Biological Materials) and then combined with primers (Integrated DNA Technologies; Table S11) and PowerUp SYBR Green Master Mix. qPCR was performed using a QuantStudio 6 Pro Real-Time PCR System and analyzed using the Design and Analysis Software version 2.4.3. Relative gene expression was calculated using the ΔCT method with *GAPDH* expression as the reference.

#### Engineered heart tissue construction

EHTs were generated in a fibrin-based hydrogel format according to the EHT Technologies product manual. Modifications were made to the composition of the hydrogel and the EHT culture media. DMEM and horse serum were replaced by RPMI 1640 basal medium, and B27 with insulin; The hydrogel was prepared by mixing the following components on ice: RPMI 1640, B27 with insulin, GlutaMAX (Thermo Fisher Scientific), water for injection, penicillin-streptomycin, ~1 mg/mL of Matrigel and 5.4 mg/mL of fibrinogen (Sigma-Aldrich). Cell mixtures were made by combining $4x10^6$ iPSC-CMs with 400 µL of hydrogel with the addition of Y-27632 (Selleck Chemicals). Polydimethylsiloxane (PDMS) racks were placed into the 2% agarose molds made in 24-well plates in preparation for EHT casting. 12 µL of 100 U/mL thrombin (Sigma-Aldrich) was added to the cell mixture immediately before casting. 100 µL of the mixture containing $1x10^6$ iPSC-CMs was transferred into the agarose mold in each well. The plate was then incubated at 37°C with 5% $CO_2$ for 1 hour to allow fibrinogen polymerization and formation of the tissue. To dissociate the EHTs from the agarose mold, 500 µL of EHT culture medium was added before shaking the plate in all directions. After a 20-minute incubation at 37°C with 5% $CO_2$, the newly formed EHTs were transferred to a new plate with 1.5 mL of medium in each well. The EHT medium consisted of RPMI 1640 with B27 supplement, 33 µg/mL of aprotinin (Sigma-Aldrich), and 50 U/mL of penicillin-streptomycin. Media changes were performed every other day and spontaneous contraction of the EHTs was observed a week post-casting.

### Supplemental Tables

**Table S1. Custom CRISPR-Cas9 oligonucleotide sequences.**

| Oligonucleotide | Sequence |
|---|---|
| IDT Alt-R® CRISPR-Cas9 crRNA | GCAAGGAATTCACTGTTACTTGG |
| IDT Ultramer® DNA Oligo HDR repair template | GGTTTGATAGCTTCCTTTAGCTCTGCAGGGCGCCC<br>AATACCAACTTGGTTTTGGGCACACACCCTGAACT<br>CATATTCCTGGTTTTCATCCAAGCTGGTAACAGTG<br>AATTCCTTGCGTACAAGCTGTG |

**Table S2. CRISPR-Cas9 on-target editing efficiencies.**

| Cell line | Genotype | Clones screened | Efficiency |
|---|---|---|---|
| AF patient iPSCs | WT/TTNtv → WT/WT | 80 | 6.3% (5/80) |

**Table S3. CRISPR-Cas9 predicted off-target sites.**

| AF patient iPSCs: WT/TTNtv → WT/WT | | | | |
|---|---|---|---|---|
| **Gene** | **Sequence** | **Mismatch** | **Locus** | **CFD Score** |
| SLC16A3 | **T**C**C**A**C**GAA**G**TCACTGTTACT GGG | 4 | chr17: 80192408-80192430:- | 0.2085 |
| KLHL2<br>KLHL2P1 | GC**C**A**A**G**G**ATTCACTGT**C**ACT GGG<br>GC**C**A**A**G**G**ATTCACTGT**C**ACT GGG | 4<br>4 | chr4: 166141149-166141171:-<br>chr4: 120255546-120255568:- | 0.1851<br>0.1851 |
| LEKR1 | GC**C**AGGAATTCACTGT**A**ACT GAG | 2 | chr3: 156763593-156763615:+ | 0.0593 |
| ELOF1 | **A**CAAGGAA**GC**CACTGTT**C**CT GGG | 4 | chr19: 11666196-11666218:+ | 0.0566 |
| LZTS3 | GCAA**A**GAATTCACTG**CC**AC**C** AGG | 4 | chr20: 3149060-3149082:+ | 0.0371 |

**Table S4. Participant characteristics at recruitment.**

| Participant | Titin genotype | Sex | Ethnicity | Age | Presentation |
|---|---|---|---|---|---|
| Early-onset AF (age 18) | Heterozygous titin truncating variant.<br>c.63624_63627delCAGC (NM_001267550.2)<br>p.Ser21134Trpfs*25 (NP_001254479.2) | Male | Asian | 56 | Paroxysmal AF managed by Sotalol.<br>Hypertension.<br>Dyslipidemia.<br>History of IgA nephropathy and mitral valve prolapse.<br>Reduced renal function.<br>Normal biventricular size, function, and geometry. |

**Table S5. Univariate linear regression of contractility measurements by genotype in beat rate-matched EHTs.**

| Cell Type | Variable | Effect Estimate | SEM | P-value |
|---|---|---|---|---|

| | | | | |
|---|---|---|---|---|
| Ventricular | Average Contraction Amplitude (mN) | 0.006942 | 0.005607 | 0.215706 |
| | Frequency (BPM) | 1.319934 | 20.87343 | 0.949579 |
| | Contraction Velocity (mN/s) | 0.020781 | 0.109776 | 0.849854 |
| | Relaxation Velocity (mN/s) | 0.015698 | 0.106628 | 0.882953 |
| | Resting Time (s) | 0.029957 | 0.054726 | 0.584099 |
| | Average Peak Widths (s) | -0.03802 | 0.06746 | 0.573025 |
| | Contraction Time T1 (s) | -0.03839 | 0.071697 | 0.592358 |
| | Relaxation Time T2 (s) | 0.00688 | 0.028481 | 0.809121 |
| Atrial | Average Contraction Amplitude (mN) | -0.01959 | 0.005727 | 0.000623 |
| | Frequency (BPM) | -8.22618 | 20.00616 | 0.680939 |
| | Contraction Velocity (mN/s) | -0.2429 | 0.082576 | 0.003266 |
| | Relaxation Velocity (mN/s) | -0.09508 | 0.045055 | 0.034831 |
| | Resting Time (s) | 0.009221 | 0.047221 | 0.845182 |
| | Average Peak Widths (s) | -0.00387 | 0.019933 | 0.846214 |
| | Contraction Time T1 (s) | 0.018951 | 0.01399 | 0.175538 |
| | Relaxation Time T2 (s) | 0.003816 | 0.033008 | 0.907958 |

**Table S6. Significantly downregulated genes in atrial WT/TTNtv relative to isogenic WT/WT EHTs.**

| Gene stable ID | Gene name | Gene description | log2 Fold Change | Adj. P-value |
|---|---|---|---|---|
| ENSG00000119411 | BSPRY | B-box and SPRY domain containing | -3.47367 | 0.019887 |
| ENSG00000137434 | C6orf52 | chromosome 6 open reading frame 52 | -3.30707 | 0.00138 |
| ENSG00000138669 | PRKG2 | protein kinase cGMP-dependent 2 | -3.26698 | 0.009476 |
| ENSG00000143171 | RXRG | retinoid X receptor gamma | -2.38815 | 0.00376 |
| ENSG00000157119 | KLHL40 | kelch like family member 40 | -4.27097 | 0.009476 |
| ENSG00000178403 | NEUROG2 | neurogenin 2 | -5.04081 | 0.040403 |
| ENSG00000184470 | TXNRD2 | thioredoxin reductase 2 | -8.04512 | 0.01302 |
| ENSG00000188321 | ZNF559 | zinc finger protein 559 | -7.552 | 0.000155 |
| ENSG00000188393 | CLEC2A | C-type lectin domain family 2 member A | -5.71134 | 1.23E-05 |
| ENSG00000206532 | RP11-553A10.1 | | -4.28411 | 2.93E-05 |
| ENSG00000217330 | SSXP10 | SSX family pseudogene 10 | -7.3713 | 0.000252 |
| ENSG00000250616 | YPEL3-DT | YPEL3 divergent transcript | -3.53372 | 0.002404 |
| ENSG00000260088 | DDX59-AS1 | DDX59 antisense RNA 1 | -4.77757 | 0.000888 |

**Table S7. Significantly upregulated genes in atrial WT/TTNtv relative to isogenic WT/WT EHTs.**

| Gene stable ID | Gene name | Gene description | log2FoldChange | Adj. P-value |
|---|---|---|---|---|
| ENSG00000053747 | LAMA3 | laminin subunit alpha 3 | 4.73246535 | 0.007761752 |
| ENSG00000075407 | ZNF37A | zinc finger protein 37A | 6.961820548 | 2.33E-10 |
| ENSG00000077943 | ITGA8 | integrin subunit alpha 8 | 4.123297475 | 0.009475613 |
| ENSG00000078081 | LAMP3 | lysosomal associated membrane protein 3 | 6.587032331 | 0.013303784 |
| ENSG00000087116 | ADAMTS2 | ADAM metallopeptidase with thrombospondin type 1 motif 2 | 4.700112636 | 0.00192069 |
| ENSG00000102802 | MEDAG | mesenteric estrogen dependent adipogenesis | 5.466979678 | 0.006310348 |
| ENSG00000106278 | PTPRZ1 | protein tyrosine phosphatase receptor type Z1 | 5.089726357 | 0.004878801 |
| ENSG00000106483 | SFRP4 | secreted frizzled related protein 4 | 6.699140575 | 0.008100616 |
| ENSG00000108846 | ABCC3 | ATP binding cassette subfamily C member 3 | 3.674480799 | 0.02443468 |
| ENSG00000109193 | SULT1E1 | sulfotransferase family 1E member 1 | 6.905738139 | 0.009475613 |
| ENSG00000112562 | SMOC2 | SPARC related modular calcium binding 2 | 5.485283399 | 0.021048328 |
| ENSG00000113209 | PCDHB5 | protocadherin beta 5 | 3.089478251 | 0.007953563 |

| | | | | |
|---|---|---|---|---|
| ENSG00000113248 | PCDHB15 | protocadherin beta 15 | 3.747232345 | 1.80E-10 |
| ENSG00000121297 | TSHZ3 | teashirt zinc finger homeobox 3 | 2.888817169 | 0.01751481 |
| ENSG00000125618 | PAX8 | paired box 8 | 7.141347937 | 0.001355237 |
| ENSG00000129317 | PUS7L | pseudouridine synthase 7 like | 8.925461098 | 1.09E-06 |
| ENSG00000130600 | H19 | H19 imprinted maternally expressed transcript | 7.264879772 | 0.03254399 |
| ENSG00000134240 | HMGCS2 | 3-hydroxy-3-methylglutaryl-CoA synthase 2 | 17.11923388 | 0.034676146 |
| ENSG00000136010 | ALDH1L2 | aldehyde dehydrogenase 1 family member L2 | 4.465388416 | 0.000137723 |
| ENSG00000140538 | NTRK3 | neurotrophic receptor tyrosine kinase 3 | 4.558324091 | 0.008206936 |
| ENSG00000145908 | ZNF300 | zinc finger protein 300 | 6.10519421 | 2.59E-07 |
| ENSG00000149451 | ADAM33 | ADAM metallopeptidase domain 33 | 4.667785584 | 0.000371649 |
| ENSG00000154277 | UCHL1 | ubiquitin C-terminal hydrolase L1 | 5.080177949 | 0.000251852 |
| ENSG00000156265 | MAP3K7CL | MAP3K7 C-terminal like | 5.50514234 | 0.001181619 |
| ENSG00000157554 | ERG | ETS transcription factor ERG | 6.687241778 | 0.012952016 |
| ENSG00000159403 | C1R | complement C1r | 3.192951937 | 0.017026672 |
| ENSG00000163661 | PTX3 | pentraxin 3 | 3.972852419 | 0.008100616 |
| ENSG00000164161 | HHIP | hedgehog interacting protein | 3.96440239 | 0.01988705 |
| ENSG00000164220 | F2RL2 | coagulation factor II thrombin receptor like 2 | 3.83713954 | 0.034445456 |
| ENSG00000165300 | SLITRK5 | SLIT and NTRK like family member 5 | 3.840579396 | 0.007137213 |
| ENSG00000166257 | SCN3B | sodium voltage-gated channel beta subunit 3 | 3.579458424 | 0.034445456 |
| ENSG00000167637 | ZNF283 | zinc finger protein 283 | 9.593265443 | 8.76E-09 |
| ENSG00000169432 | SCN9A | sodium voltage-gated channel alpha subunit 9 | 3.380581054 | 0.000616209 |
| ENSG00000170477 | KRT4 | keratin 4 | 5.592746472 | 0.02443468 |
| ENSG00000171291 | ZNF439 | zinc finger protein 439 | 5.503237958 | 6.21E-14 |
| ENSG00000171466 | ZNF562 | zinc finger protein 562 | 8.846749211 | 5.22E-17 |
| ENSG00000174080 | CTSF | cathepsin F | 5.722741884 | 0.007953563 |
| ENSG00000176971 | FIBIN | fin bud initiation factor homolog | 4.252020748 | 0.005431214 |
| ENSG00000177932 | ZNF354C | zinc finger protein 354C | 3.85601352 | 0.030869795 |
| ENSG00000178187 | ZNF454 | zinc finger protein 454 | 4.465514068 | 0.005243655 |
| ENSG00000179542 | SLITRK4 | SLIT and NTRK like family member 4 | 2.884967354 | 0.022471867 |
| ENSG00000180543 | TSPYL5 | TSPY like 5 | 12.67580153 | 2.12E-16 |
| ENSG00000181291 | TMEM132E | transmembrane protein 132E | 4.324064453 | 0.002848939 |

| | | | | |
|---|---|---|---|---|
| ENSG00000183715 | OPCML | opioid binding protein/cell adhesion molecule like | 2.548859592 | 0.038679978 |
| ENSG00000185070 | FLRT2 | fibronectin leucine rich transmembrane protein 2 | 3.224744513 | 0.015020896 |
| ENSG00000185215 | TNFAIP2 | TNF alpha induced protein 2 | 4.060394089 | 0.02932957 |
| ENSG00000185869 | ZNF829 | zinc finger protein 829 | 9.512833358 | 2.33E-08 |
| ENSG00000186446 | ZNF501 | zinc finger protein 501 | 5.16042094 | 2.00E-07 |
| ENSG00000187955 | COL14A1 | collagen type XIV alpha 1 chain | 3.711935178 | 1.46E-05 |
| ENSG00000187957 | DNER | delta/notch like EGF repeat containing | 4.419727036 | 0.029458148 |
| ENSG00000189129 | PLAC9 | placenta associated 9 | 4.650679649 | 0.04997267 |
| ENSG00000189223 | PAX8-AS1 | PAX8 antisense RNA 1 | 8.844898287 | 1.65E-19 |
| ENSG00000196268 | ZNF493 | zinc finger protein 493 | 4.32394697 | 0.000222539 |
| ENSG00000196867 | ZFP28 | ZFP28 zinc finger protein | 8.3601092 | 3.62E-08 |
| ENSG00000197608 | ZNF841 | zinc finger protein 841 | 6.717254931 | 0.00583753 |
| ENSG00000197928 | ZNF677 | zinc finger protein 677 | 11.58725394 | 7.03E-13 |
| ENSG00000198001 | IRAK4 | interleukin 1 receptor associated kinase 4 | 8.054837003 | 1.52E-05 |
| ENSG00000198121 | LPAR1 | lysophosphatidic acid receptor 1 | 3.525742039 | 0.001355237 |
| ENSG00000198768 | APCDD1L | APC down-regulated 1 like | 7.477195725 | 0.008579337 |
| ENSG00000204969 | PCDHA2 | protocadherin alpha 2 | 6.280341079 | 0.021734506 |
| ENSG00000205038 | PKHD1L1 | PKHD1 like 1 | 8.223526111 | 0.022373684 |
| ENSG00000205403 | CFI | complement factor I | 3.446871213 | 0.02443468 |
| ENSG00000215146 | | zinc finger protein pseudogene | 8.131461525 | 7.54E-05 |
| ENSG00000224597 | SVIL-AS1 | SVIL antisense RNA 1 | 6.634598033 | 0.010415631 |
| ENSG00000227124 | ZNF717 | zinc finger protein 717 | 8.089604774 | 7.48E-09 |
| ENSG00000234773 | | zinc finger protein pseudogene | 2.536211107 | 0.042637422 |
| ENSG00000245680 | ZNF585B | zinc finger protein 585B | 2.694229782 | 0.005666615 |
| ENSG00000247516 | MIR4458HG | MIR4458 host gene | 7.986597856 | 3.46E-05 |
| ENSG00000250120 | PCDHA10 | protocadherin alpha 10 | 2.954184065 | 1.66E-05 |
| ENSG00000253731 | PCDHGA6 | protocadherin gamma subfamily A, 6 | 3.620141284 | 0.031414742 |
| ENSG00000253767 | PCDHGA8 | protocadherin gamma subfamily A, 8 | 2.803143838 | 0.003298428 |
| ENSG00000253953 | PCDHGB4 | protocadherin gamma subfamily B, 4 | 2.543059282 | 3.46E-05 |
| ENSG00000254221 | PCDHGB1 | protocadherin gamma subfamily B, 1 | 3.963078142 | 0.022373684 |
| ENSG00000255366 | | novel lncRNA | 7.478968041 | 0.004854819 |

| | | | | |
|---|---|---|---|---|
| ENSG00000263711 | LINC02864 | long intergenic non-protein coding RNA 2864 | 9.750600837 | 1.18E-08 |
| ENSG00000267254 | ZNF790-AS1 | ZNF790 antisense RNA 1 | 5.158361744 | 0.03254399 |
| ENSG00000267454 | ZNF582-DT | ZNF582 divergent transcript | 4.496082871 | 0.036655923 |
| ENSG00000278318 | ZNF229 | zinc finger protein 229 | 8.066497713 | 3.67E-12 |
| ENSG00000278897 | | | 6.914451586 | 0.000251852 |
| ENSG00000281406 | BLACAT1 | BLACAT1 overlapping LEMD1 locus | 6.811191735 | 0.012952016 |

**Table S8. Significantly upregulated genes in ventricular WT/TTNtv relative to isogenic WT/WT EHTs.**

| Gene stable ID | Gene name | Gene description | log2FoldChange | Adj. P-value |
|---|---|---|---|---|
| ENSG00000075407 | ZNF37A | zinc finger protein 37A | 7.553598887 | 6.21E-08 |
| ENSG00000110446 | SLC15A3 | solute carrier family 15 member 3 | 4.677662572 | 0.037842 |
| ENSG00000113248 | PCDHB15 | protocadherin beta 15 | 3.483071755 | 6.27E-09 |
| ENSG00000119917 | IFIT3 | interferon induced protein with tetratricopeptide repeats 3 | 4.585193712 | 0.000108 |
| ENSG00000120322 | PCDHB8 | protocadherin beta 8 | 2.750884908 | 0.014655 |
| ENSG00000125618 | PAX8 | paired box 8 | 5.273166136 | 0.03645 |
| ENSG00000129317 | PUS7L | pseudouridine synthase 7 like | 7.172293317 | 3.08E-07 |
| ENSG00000130600 | H19 | H19 imprinted maternally expressed transcript | 8.113800368 | 0.0118 |
| ENSG00000134321 | RSAD2 | radical S-adenosyl methionine domain containing 2 | 4.487811019 | 0.007546 |
| ENSG00000136010 | ALDH1L2 | aldehyde dehydrogenase 1 family member L2 | 3.910717703 | 0.006694 |
| ENSG00000145908 | ZNF300 | zinc finger protein 300 | 6.077428648 | 3.41E-06 |
| ENSG00000148053 | NTRK2 | neurotrophic receptor tyrosine kinase 2 | 2.422081661 | 0.005305 |
| ENSG00000167637 | ZNF283 | zinc finger protein 283 | 6.113981747 | 4.87E-06 |
| ENSG00000171291 | ZNF439 | zinc finger protein 439 | 5.498734294 | 2.07E-11 |
| ENSG00000171466 | ZNF562 | zinc finger protein 562 | 7.501896608 | 9.13E-14 |
| ENSG00000174080 | CTSF | cathepsin F | 6.363698718 | 0.049949 |
| ENSG00000177932 | ZNF354C | zinc finger protein 354C | 4.662710977 | 0.004087 |
| ENSG00000180543 | TSPYL5 | TSPY like 5 | 8.426718801 | 1.45E-24 |
| ENSG00000185869 | ZNF829 | zinc finger protein 829 | 6.801090794 | 4.45E-05 |
| ENSG00000186446 | ZNF501 | zinc finger protein 501 | 5.519309359 | 3.41E-06 |
| ENSG00000188313 | PLSCR1 | phospholipid scramblase 1 | 3.172239557 | 0.016319 |
| ENSG00000189223 | PAX8-AS1 | PAX8 antisense RNA 1 | 9.367161469 | 9.13E-14 |

| | | | | |
|---|---|---|---|---|
| ENSG00000196268 | ZNF493 | zinc finger protein 493 | 4.514693814 | 0.000164 |
| ENSG00000196867 | ZFP28 | ZFP28 zinc finger protein | 7.655048369 | 8.78E-09 |
| ENSG00000197928 | ZNF677 | zinc finger protein 677 | 7.863501949 | 5.55E-12 |
| ENSG00000198001 | IRAK4 | interleukin 1 receptor associated kinase 4 | 8.56279668 | 2.20E-06 |
| ENSG00000215146 | | zinc finger protein pseudogene | 6.836732629 | 0.000444 |
| ENSG00000227124 | ZNF717 | zinc finger protein 717 | 5.727448646 | 4.06E-06 |
| ENSG00000247516 | MIR4458HG | MIR4458 host gene | 6.189238115 | 0.010693 |
| ENSG00000253953 | PCDHGB4 | protocadherin gamma subfamily B, 4 | 2.537513048 | 6.33E-05 |
| ENSG00000254221 | PCDHGB1 | protocadherin gamma subfamily B, 1 | 5.616136067 | 0.008617 |
| ENSG00000263711 | LINC02864 | long intergenic non-protein coding RNA 2864 | 8.478985464 | 1.31E-06 |
| ENSG00000272674 | PCDHB16 | protocadherin beta 16 | 2.423355024 | 0.014768 |
| ENSG00000278318 | ZNF229 | zinc finger protein 229 | 6.571400241 | 6.59E-10 |
| ENSG00000278897 | | | 5.761487035 | 0.003287 |

**Table S9. Significantly differentially spliced genes in atrial WT/TTNtv and isogenic WT/WT EHTs.**

| Gene Name | Exon Start | Exon End | Ratio difference | Adj. P-value | Splicing event type |
|---|---|---|---|---|---|
| COL16A1 | 31702120 | 31702227 | -0.303061928 | 0.011207 | Spliced Exon |
| TPM1 | 63061197 | 63061273 | -0.170830725 | 4.49E-14 | Spliced Exon |
| SIPA1L2 | 232533489 | 232533583 | -0.119322096 | 0.032502 | Spliced Exon |
| SIPA1L2 | 232574173 | 232574222 | -0.11464023 | 0.021224 | Spliced Exon |
| FHL2 | 105396646 | 105396697 | -0.100184577 | 4.32E-06 | Spliced Exon |
| AC109583.1 | 46811105 | 46811873 | -0.202607315 | 1.07E-22 | Spliced Exon |
| MAP4 | 47909037 | 47912421 | 0.129839162 | 0.016973 | Spliced Exon |
| RBM24 | 17290848 | 17290893 | 0.107057492 | 0.005013 | Spliced Exon |
| LAMA2 | 129475389 | 129475401 | 0.147056086 | 0.026181 | Spliced Exon |
| ACTB | 5529018 | 5529059 | 0.120249052 | 1.08E-16 | Spliced Exon |
| ATP5F1C | 7806973 | 7807010 | -0.220128247 | 4.18E-06 | Spliced Exon |
| POSTN | 37574571 | 37574652 | 0.297028903 | 0.005883 | Spliced Exon |
| ACTN1 | 68878988 | 68879069 | -0.206702849 | 0.001749 | Spliced Exon |
| ACTG1 | 81511762 | 81511803 | 0.231863818 | 3.25E-58 | Spliced Exon |
| TMEM259 | 1010990 | 1011095 | 0.125036128 | 0.013237 | Spliced Exon |

| | | | | | |
|---|---|---|---|---|---|
| EMC10 | 50481860 | 50481947 | 0.127787939 | 6.10E-12 | Spliced Exon |
| GNAS | 58889525 | 58889549 | -0.209209106 | 2.39E-16 | Spliced Exon |
| TNRC6B | 40281118 | 40281289 | 0.388665087 | 0.003459 | Spliced Exon |
| AC109583.1 | 46811105 | 46811873 | -0.202607315 | 5.16E-23 | Retained Intron |
| TPM1 | 63061197 | 63061273 | -0.170830725 | 3.81E-14 | Mutually Exclusive Exon |
| ERBB4 | 211657753 | 211657828 | -0.270791406 | 0.010619 | Mutually Exclusive Exon |
| RBM24 | 17290848 | 17290893 | 0.107057492 | 0.005059 | Mutually Exclusive Exon |
| ACTN1 | 68878988 | 68879069 | -0.206702849 | 0.001695 | Mutually Exclusive Exon |
| TMEM259 | 1010990 | 1011095 | 0.125036128 | 0.011756 | Mutually Exclusive Exon |
| MIB2 | 1624964 | 1624989 | -0.241959315 | 0.04038 | Alternative 3' Splice Site |
| TPM1 | 63043468 | 63043704 | -0.149459796 | 2.10E-13 | Alternative 3' Splice Site |
| TPM1 | 63061197 | 63061711 | -0.145303259 | 2.38E-10 | Alternative 3' Splice Site |
| PDLIM5 | 94575615 | 94575941 | -0.151059192 | 0.000354 | Alternative 3' Splice Site |
| PDE4DIP | 148992254 | 148993483 | 0.216695568 | 0.010971 | Alternative 3' Splice Site |
| MYL7 | 44140428 | 44140517 | -0.109655235 | 1.17E-116 | Alternative 3' Splice Site |
| KCP | 128891088 | 128891096 | -0.617695153 | 0.008604 | Alternative 3' Splice Site |
| B4GALNT3 | 545139 | 545367 | 0.202645055 | 0.000134 | Alternative 3' Splice Site |
| ENO2 | 6915435 | 6915819 | -0.107052514 | 1.65E-05 | Alternative 3' Splice Site |
| ENO2 | 6915435 | 6915816 | -0.107585547 | 1.62E-05 | Alternative 3' Splice Site |
| KRT8 | 52949579 | 52949585 | -0.101549952 | 0.047282 | Alternative 3' Splice Site |
| SERF2 | 43798868 | 43798942 | 0.246993566 | 0.0286 | Alternative 3' Splice Site |
| HCFC1R1 | 3023531 | 3023968 | 0.256674858 | 1.93E-05 | Alternative 3' Splice Site |
| ACTG1 | 81511627 | 81511803 | 0.18912013 | 1.64E-36 | Alternative 3' Splice Site |
| TPM1 | 63060940 | 63061273 | -0.144331992 | 3.26E-10 | Alternative 5' Splice Site |
| MYL3 | 46861296 | 46861684 | -0.114943173 | 2.55E-25 | Alternative 5' Splice Site |
| CRYAB | 111913596 | 111913897 | 0.277299771 | 6.40E-16 | Alternative 5' Splice Site |
| ATP2A2 | 110346322 | 110347791 | -0.102499935 | 4.02E-18 | Alternative 5' Splice Site |
| YPEL3 | 30096614 | 30096915 | 0.314238159 | 0.000203 | Alternative 5' Splice Site |
| ZFP36 | 39406929 | 39407109 | -0.212587558 | 0.003216 | Alternative 5' Splice Site |
| GNAS | 58889214 | 58889353 | -0.194003668 | 1.37E-14 | Alternative 5' Splice Site |
| GNAS | 58889333 | 58889353 | -0.177813552 | 3.85E-14 | Alternative 5' Splice Site |

| | | | | | |
|---|---|---|---|---|---|
| GNAS | 58889354 | 58889549 | -0.202198528 | 6.40E-16 | Alternative 5' Splice Site |
| GNAS | 58889214 | 58889549 | -0.200449243 | 1.00E-15 | Alternative 5' Splice Site |
| GNAS | 58889407 | 58889549 | -0.20552399 | 9.81E-17 | Alternative 5' Splice Site |
| GNAS | 58889333 | 58889549 | -0.200125858 | 8.41E-16 | Alternative 5' Splice Site |
| GNAS | 58889354 | 58889406 | -0.185899016 | 9.04E-13 | Alternative 5' Splice Site |
| GNAS | 58889214 | 58889406 | -0.192896132 | 3.04E-14 | Alternative 5' Splice Site |
| GNAS | 58889333 | 58889406 | -0.185746182 | 2.99E-13 | Alternative 5' Splice Site |
| GNAS | 58889214 | 58889332 | -0.19616702 | 1.26E-14 | Alternative 5' Splice Site |

**Table S10. Significantly differentially spliced genes in ventricular WT/TTNtv and isogenic WT/WT EHTs.**

| Gene Name | Exon Start | Exon End | Ratio difference | Adj. P-value | Splicing event type |
|---|---|---|---|---|---|
| ADCY5 | 123296979 | 123297054 | 0.13876064 | 2.04E-07 | Spliced Exon |
| RPS24 | 78040200 | 78040225 | 0.152983148 | 0.003475 | Spliced Exon |
| RPS24 | 78037964 | 78037982 | 0.201641219 | 3.13E-05 | Spliced Exon |
| RPS24 | 78040203 | 78040225 | 0.145676338 | 0.002271 | Spliced Exon |
| RPS24 | 78037964 | 78037993 | 0.216746383 | 0.001785 | Spliced Exon |
| NACA | 56715870 | 56716110 | 0.22077717 | 0.002591 | Spliced Exon |
| WARS1 | 100375282 | 100375406 | -0.314919305 | 0.042279 | Spliced Exon |
| WARS1 | 100375282 | 100375403 | -0.313966667 | 0.042279 | Spliced Exon |
| ACTG1 | 81511762 | 81511803 | 0.166542068 | 4.30E-11 | Spliced Exon |
| TPM4 | 16095263 | 16095357 | 0.309618907 | 0.003455 | Spliced Exon |
| GNAS | 58889525 | 58889549 | -0.151298113 | 1.72E-08 | Spliced Exon |
| EIF4G2 | 10808704 | 10808940 | 0.134168855 | 0.000783 | Retained Intron |
| RPS24 | 78037964 | 78037982 | 0.201641219 | 4.30E-05 | Mutually Exclusive Exon |
| RPS24 | 78037964 | 78037993 | 0.216746383 | 0.001358 | Mutually Exclusive Exon |
| NACA | 56715870 | 56716110 | 0.22077717 | 0.001882 | Mutually Exclusive Exon |
| WARS1 | 100375282 | 100375406 | -0.314919305 | 0.027557 | Mutually Exclusive Exon |
| WARS1 | 100375282 | 100375350 | -0.291749779 | 0.043183 | Mutually Exclusive Exon |
| WARS1 | 100375282 | 100375403 | -0.313966667 | 0.027557 | Mutually Exclusive Exon |
| TNNT2 | 201367804 | 201367806 | 0.104067197 | 6.03E-14 | Alternative 3' Splice Site |
| MAP4 | 47852939 | 47853039 | -0.158068226 | 0.012278 | Alternative 3' Splice Site |

| | | | | | |
|---|---|---|---|---|---|
| VDAC1 | 133993016 | 133993018 | -0.127367196 | 0.00014 | Alternative 3' Splice Site |
| WARS1 | 100375351 | 100375406 | -0.340871859 | 0.03057 | Alternative 3' Splice Site |
| WARS1 | 100375351 | 100375403 | -0.340314706 | 0.03057 | Alternative 3' Splice Site |
| CTDNEP1 | 7251413 | 7251601 | 0.203556757 | 4.04E-07 | Alternative 3' Splice Site |
| ACTG1 | 81511627 | 81511803 | 0.12124877 | 0.000142 | Alternative 3' Splice Site |
| HSPB1 | 76303845 | 76303888 | 0.109949247 | 5.50E-29 | Alternative 5' Splice Site |
| HSPB1 | 76303845 | 76303882 | 0.10864134 | 4.37E-31 | Alternative 5' Splice Site |
| CRYAB | 111910632 | 111910852 | -0.107059692 | 0.001672 | Alternative 5' Splice Site |
| CRYAB | 111910632 | 111910830 | -0.111446648 | 0.001015 | Alternative 5' Splice Site |
| CRYAB | 111910669 | 111910852 | -0.105345815 | 0.002644 | Alternative 5' Splice Site |
| GNAS | 58889214 | 58889353 | -0.219466034 | 9.08E-19 | Alternative 5' Splice Site |
| GNAS | 58889333 | 58889353 | -0.229812597 | 2.02E-19 | Alternative 5' Splice Site |
| GNAS | 58889354 | 58889549 | -0.164079373 | 2.12E-10 | Alternative 5' Splice Site |
| GNAS | 58889214 | 58889549 | -0.18843914 | 5.73E-14 | Alternative 5' Splice Site |
| GNAS | 58889407 | 58889549 | -0.171590295 | 3.70E-11 | Alternative 5' Splice Site |
| GNAS | 58889333 | 58889549 | -0.172845968 | 2.02E-11 | Alternative 5' Splice Site |
| GNAS | 58889354 | 58889406 | -0.140911576 | 5.71E-08 | Alternative 5' Splice Site |
| GNAS | 58889214 | 58889406 | -0.201006346 | 5.23E-16 | Alternative 5' Splice Site |
| GNAS | 58889333 | 58889406 | -0.175522242 | 9.36E-12 | Alternative 5' Splice Site |
| GNAS | 58889214 | 58889332 | -0.21646305 | 1.05E-18 | Alternative 5' Splice Site |

**Table S11. Primer sequences.**

| Assay | Gene/Target | Sequence (5'-3') |
|---|---|---|
| PCR: genotyping by Sanger sequencing | TTNtv mutation | TACAGAATCTCCTCCCAGACCA |
| | | GCTCTCACTATGACCAGTTTCC |
| | SLC16A3 | TACCAGTATGAGCGGGACCT |
| | | CCCCTTTCCTTGGGTTCCTC |
| | KLHL2 KLHL2P1 | GAACTTGGACACCAGGGAGG |
| | | ATAGAGGAGCCCCACACCAT |
| | LEKR1 | CCACCACTTTCCCAACCTCA |
| | | ATCTGCTTGTAGTGGCGCAA |
| | ELOF1 | AGAGGAGACTTGTGGGGCT |
| | | CCGCAGTTCCTCAAAAGCAC |
| | LZTS3 | ACTGACGGAGCCTCTCTCAG |
| | | TCCCTGCCTGGTTTATGCAG |
| RT-qPCR: iPSC-CM characterization | TNNT2 | TTCACCAAAGATCTGCTCCT |
| | | TACTGGTGTGGAGTGGGTG |
| | TTN All | GTAAAAAGAGCTGCCCCAGTGA |
| | | GCTAGGTGGCCCAGTGCTACT |
| | IRX4 | TTCCGTTCTGAAGCGTGGTC |
| | | TGAAGCAGGCAATTATTGGTGT |
| | MYL2 | ACAGGGATGGCTTCATTGAC |
| | | CCGCTCCCTTAAGTTTCTCC |
| | TBX5 | CTTGTGATGTTTTCAGAGCC |
| | | TTCTCTCTAAAAGCAAGCGT |
| | NPPA | ACAGGATTGGAGCCCAGAG |
| | | GGAGCCTCTTGCAGTCTGTC |
| | KCNJ3 | CTGCTCAAAGGATGACTTGT |
| | | CATGGAACTGGGAGTAATCA |
| | KCNA5 | CGAGGATGAGGGCTTCATTA |
| | | CTGAACTCAGGCAGGGTCTC |
| | GAPDH | CAAGAGCACAAGAGGAAGAGAG |
| | | CTACATGGCAACTGTGAGGAG |

### Supplemental Figures

**Figure S1. Atrial and ventricular iPSC-CM characterization.** **A**, Heatmap showing expression of cell type-specific markers in atrial and ventricular iPSC-CMs (n = 3 per cell type; n = 3 per genotype). Colours represent z-scores of normalized gene counts. Comparisons of **B**, beat rate, and **C**, rate-corrected action potential at 30% repolarization in WT/TTNtv iPSC-CMs versus isogenic WT/WT controls. TTNtv, titin truncating variant; WT, wild-type; t-test.

**Figure S2. Atrial and ventricular EHT characterization.** **A**, Heatmap showing expression of cell type-specific markers in atrial and ventricular EHT samples (n = 3 per cell type; n = 3 per genotype). Colours represent z-scores of normalized gene counts. **B**, Diagram of contractility variables measured. **C**, Contraction (T1) and relaxation (T2) times of atrial EHTs (n = 8-17 EHTs). **D**, Contraction and relaxation velocities of atrial EHTs (n = 8-17 EHTs). **E**, Contraction and relaxation velocities of ventricular EHTs (n = 10-17 EHTs). **F**, Contraction (T1) and relaxation (T2) times of ventricular EHTs (n = 10-17 EHTs). **G**, Spontaneous beat rates of atrial and ventricular EHTs (n = 8-17 EHTs). EHT indicates engineered heart tissue; TTNtv, titin truncating variant; WT, wild-type; Vel., velocity; $*P < 0.05$; $**P < 0.01$; t-test.

**Figure S3. Differential splicing analysis of atrial EHTs.** Volcano plots of genes with **A**, skipped exons, **B**, mutually excluded exons, **C**, retained introns, **D**, alternative 5’ splice sites, and **E**, alternative 3’ splice sites in atrial WT/TTNtv and WT/WT EHTs (ratio difference < 0 indicates a splicing event in WT/WT EHTs; ratio difference > 0 indicates an event in WT/TTNtv EHTs). Events with an FDR-adjusted *P*-value < 0.05 and absolute ratio difference > 0.1 are considered significant. EHT indicates engineered heart tissue; TTNtv, titin truncating variant; WT, wild-type.

**Figure S4. Differential splicing analysis of ventricular EHTs.** Volcano plots of genes with **A**, skipped exons, **B**, mutually excluded exons, **C**, retained introns, **D**, alternative 5’ splice sites, and **E**, alternative 3’ splice sites in ventricular WT/TTNtv and WT/WT EHTs (ratio difference < 0 indicates a splicing event in WT/WT EHTs; ratio difference > 0 indicates an event in WT/TTNtv EHTs). Events with an FDR-adjusted *P*-value < 0.05 and absolute ratio difference > 0.1 are considered significant. EHT indicates engineered heart tissue; TTNtv, titin truncating variant; WT, wild-type.
