## Supplementary figures and images for "Patient and cell-type specific hiPSC-modeling of a truncating titin variant associated with atrial fibrillation"

### Supplemental Figures

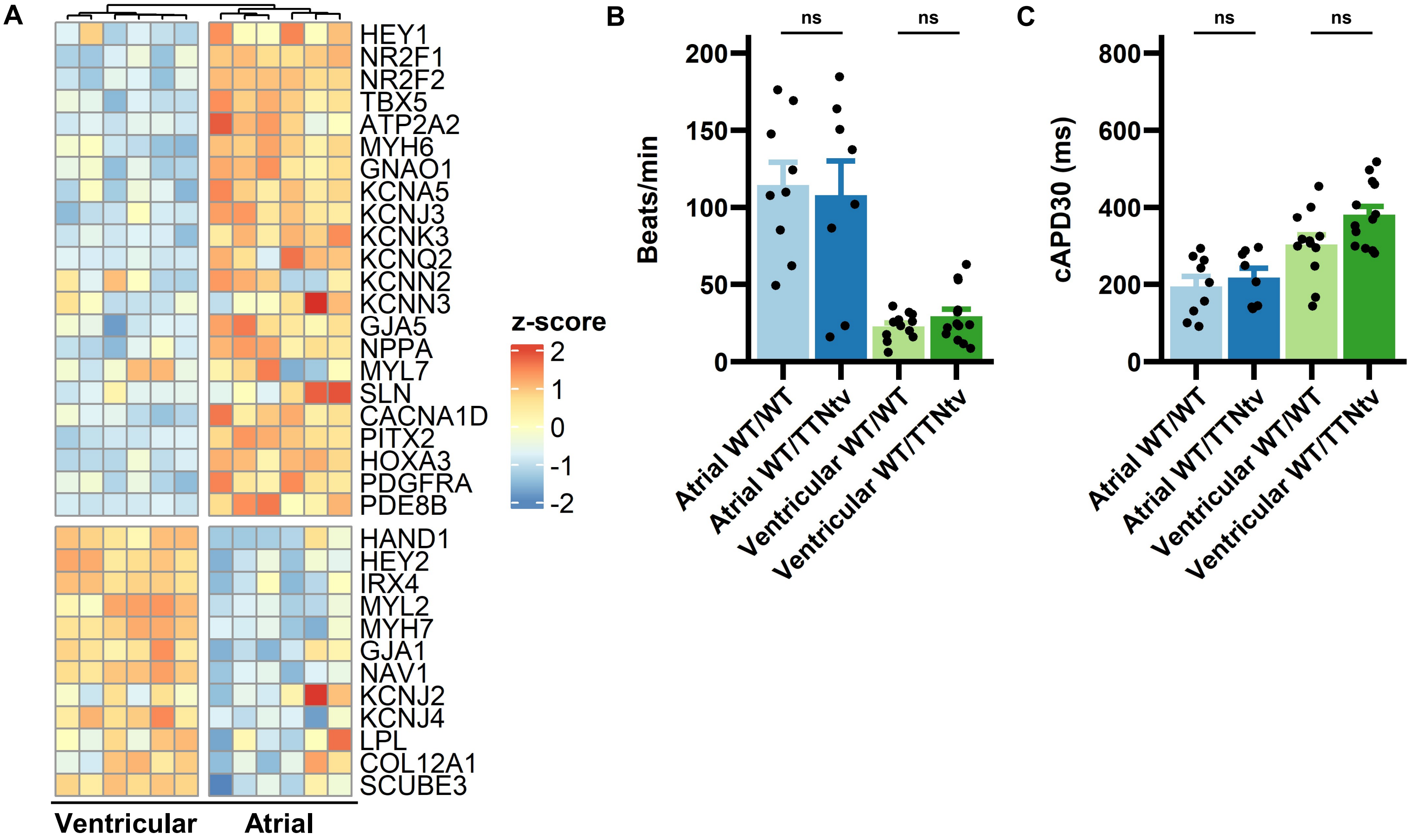

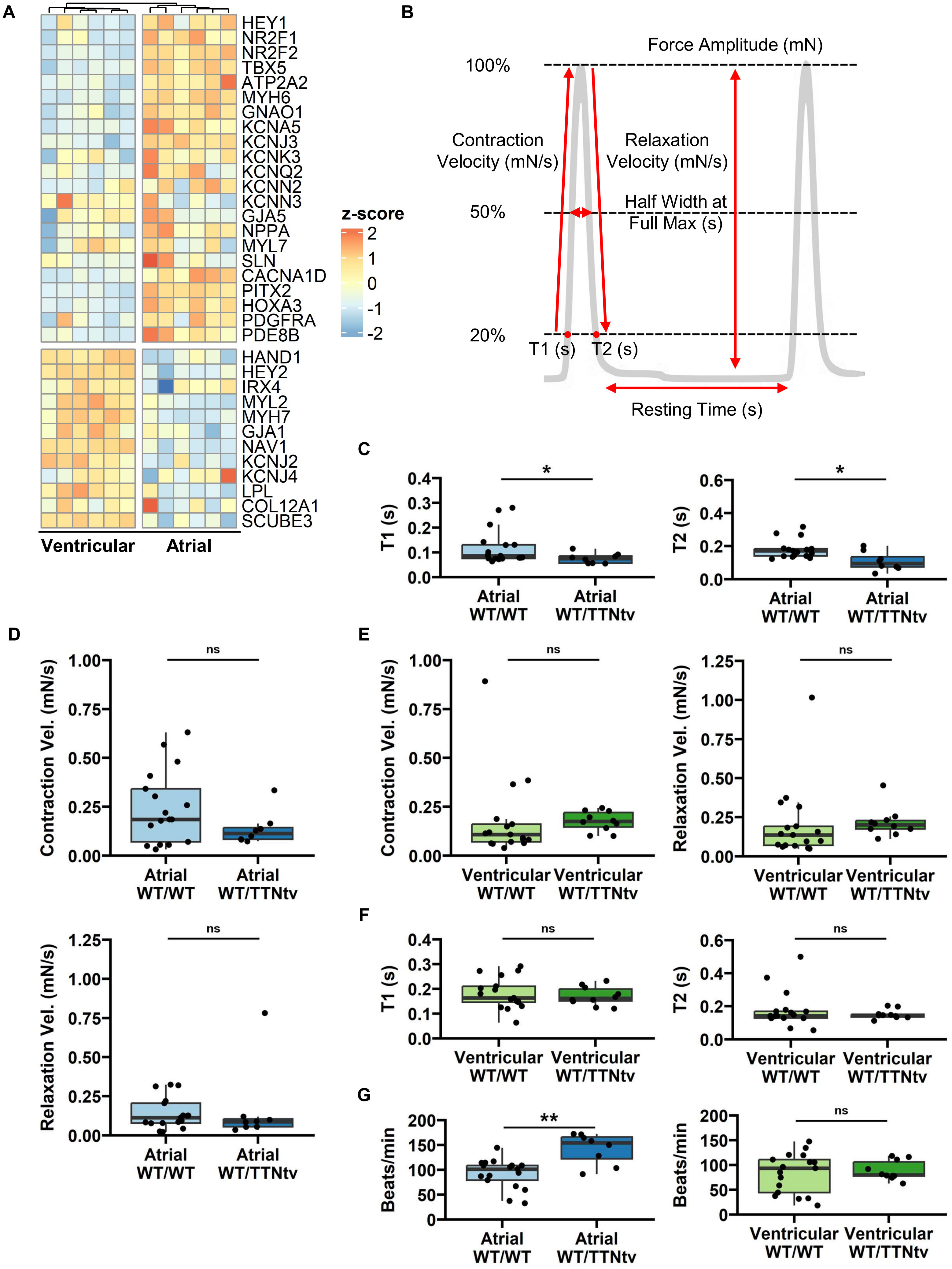

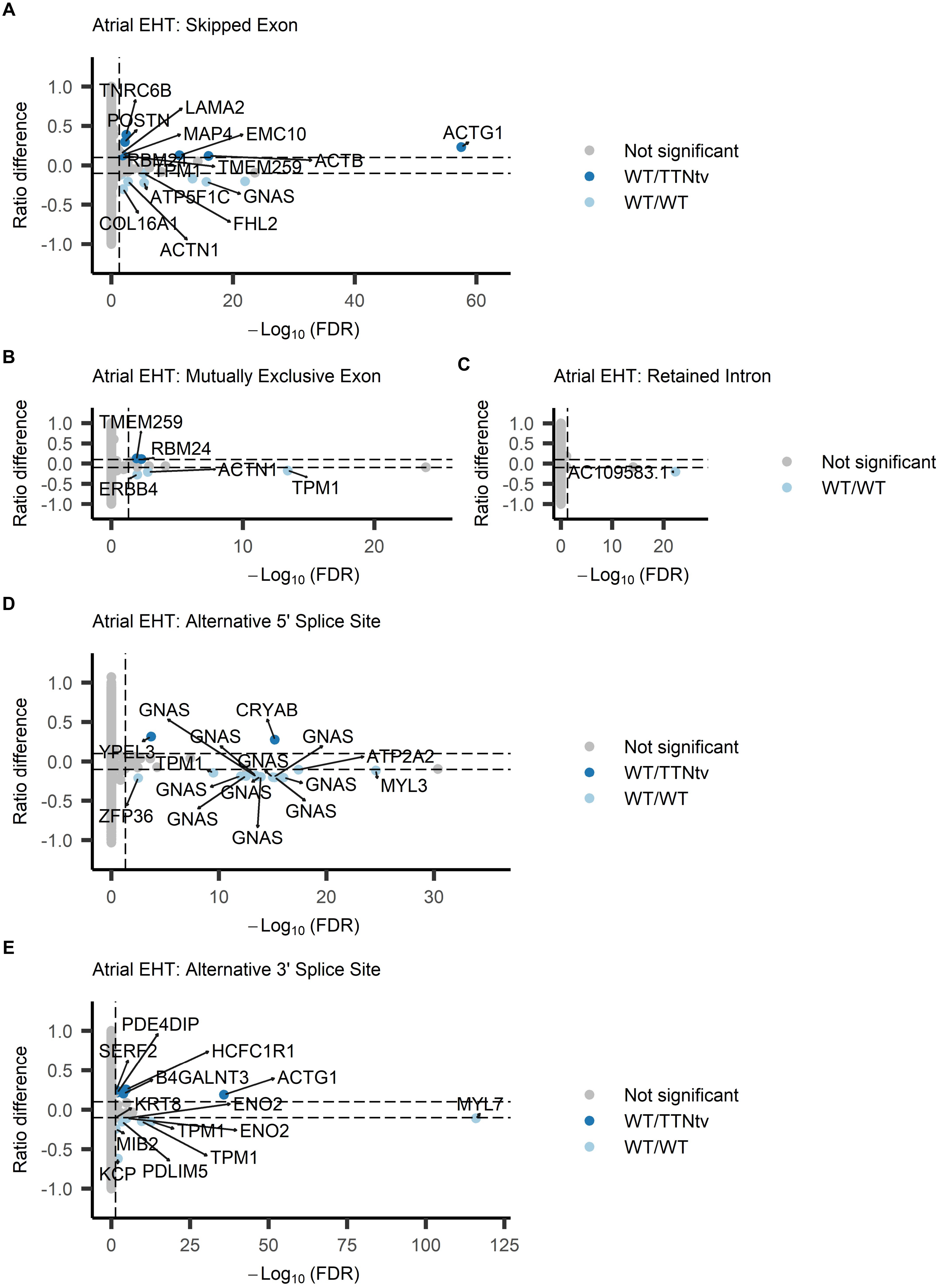

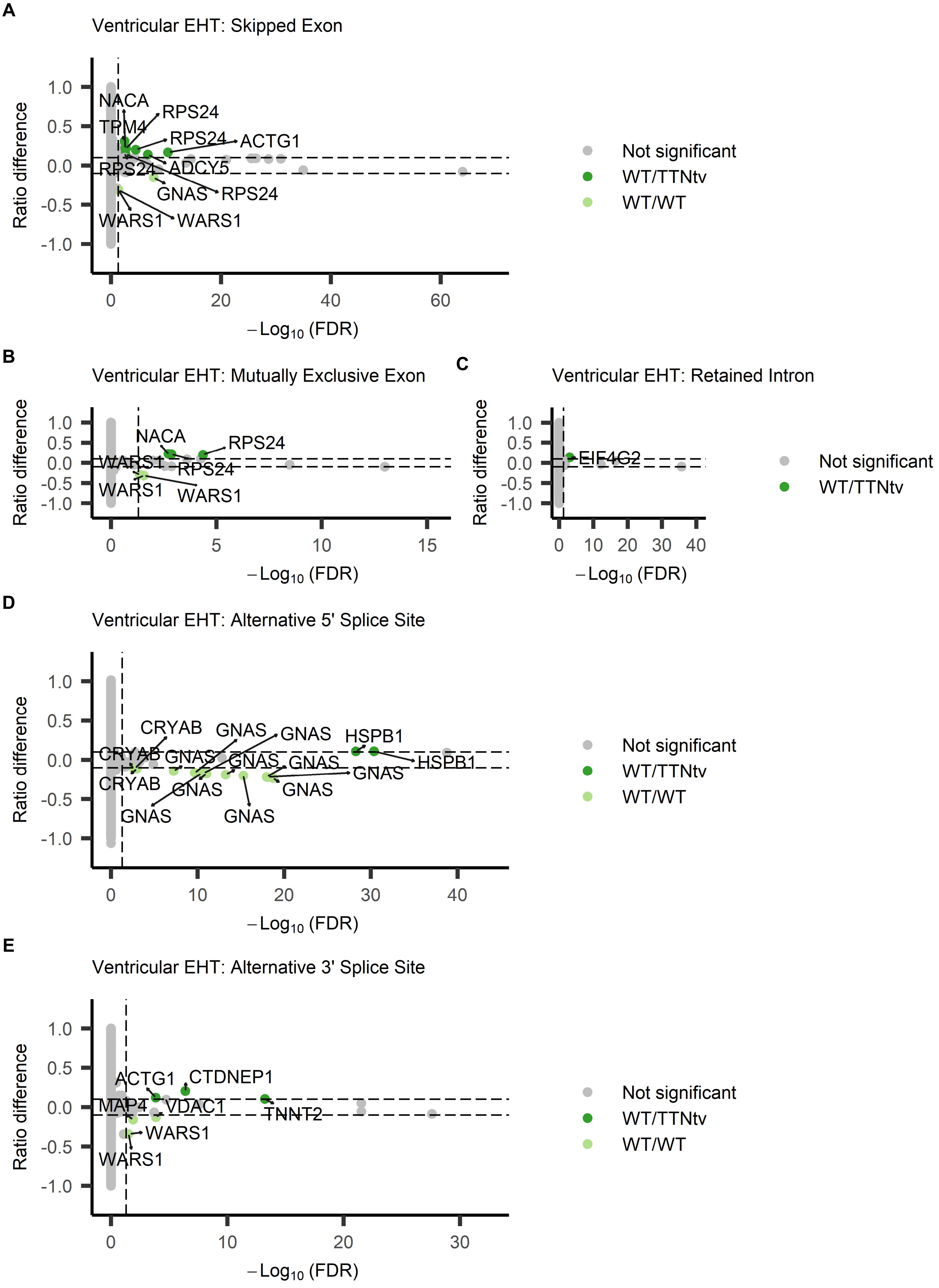
